# Comprehensive benchmarking of computational deconvolution of transcriptomics data

**DOI:** 10.1101/2020.01.10.897116

**Authors:** Francisco Avila Cobos, José Alquicira-Hernandez, Joseph Powell, Pieter Mestdagh, Katleen De Preter

**Author notes:** These authors contributed equally to this work (Joseph Powell, Pieter Mestdagh, Katleen De Preter).

## Abstract

Many computational methods to infer cell type proportions from bulk transcriptomics data have been developed. Attempts comparing these methods revealed that the choice of reference marker signatures is far more important than the method itself. However, a thorough evaluation of the combined impact of data transformation, pre-processing, marker selection, cell type composition and choice of methodology on the results is still lacking.

Using different single-cell RNA-sequencing (scRNA-seq) datasets, we generated hundreds of pseudo-bulk mixtures to evaluate the combined impact of these factors on the deconvolution results. Along with methods to perform deconvolution of bulk RNA-seq data we also included five methods specifically designed to infer the cell type composition of bulk data using scRNA-seq data as reference.

Both bulk and single-cell deconvolution methods perform best when applied to data in linear scale and the choice of normalization can have a dramatic impact on the performance of some, but not all methods. Overall, single-cell methods have comparable performance to the best performing bulk methods and bulk methods based on semi-supervised approaches showed higher error and lower correlation values between the computed and the expected proportions. Moreover, failure to include cell types in the reference that are present in a mixture always led to substantially worse results, regardless of any of the previous choices. Taken together, we provide a thorough evaluation of the combined impact of the different factors affecting the computational deconvolution task across different datasets and propose general guidelines to maximize its performance.

## Introduction

Since bulk samples of heterogeneous mixtures only represent averaged expression levels (rather than individual measures for each gene across different cell types present in such mixture), many relevant analyses such as differential gene expression are typically confounded by differences in cell type proportions. Moreover, understanding differences in cell type composition in diseases such as cancer may enable scientists to identify potentially interesting cellular populations to be targeted therapeutically. For instance, the abundance of tumor infiltrating lymphocytes and other immune cells in solid tumors (also known as the tumor microenvironment) is currently a very active field of research^1–3^ (e.g. in the context of immunotherapy) and it has already been shown that accounting for the tumor heterogeneity resulted in more sensitive survival analyses and more accurate tumor subtype predictions^4^. For these reasons, many methodologies to infer proportions of individual cell types (= computational deconvolution) from bulk transcriptomics data have been developed during the last two decades^5^ and various methods able to use single-cell RNA-sequencing data have emerged in the past year alone.

Several studies have addressed different factors affecting the deconvolution results but only focused on one or two individual aspects at a time. For instance, Zhong and Liu^6^ showed that applying the logarithmic transformation to microarray data led to a consistent under-estimation of cell-type specific expression profiles. Hoffmann *et al*.^7^ showed that four different normalization strategies had an impact on the estimation of cell type proportions from microarray data and Newman *et al*.^8^ highlighted the importance of accounting for differences in normalization procedures when comparing the results from CIBERSORT^9^ and TIMER^10^. Furthermore, Vallania *et al*.^11^ observed highly concordant results across different deconvolution methods in both blood and tissue samples, suggesting that the reference matrix was more important than the methodology being used.

Sturm *et al*.^12^ already investigated scenarios where reported cell type proportions were higher than expected (spillover effect) or different from zero when a cell type was not present in a mixture (background prediction), possibly caused by related cell types sharing similar signatures or marker genes not being sufficiently cell-type specific. Moreover, they provided a guideline for method selection depending on which cell type of interest needs to be deconvolved. However, each method evaluated in Sturm *et al.* was accompanied by its own reference signature for the different immune cell types, implying that differences may be marker-dependent and not method-dependent. Moreover, they did not evaluate the effect of data transformation and normalization in these analyses and only focused on immune cell types.

Here we provide a comprehensive and quantitative evaluation of the combined impact of data transformation, scaling/normalization, marker selection, cell type composition and choice of methodology on the deconvolution results. In this study we evaluated the performance of 20 deconvolution methods aimed at computing cell type proportions, including five recently developed methods that use single-cell RNA-sequencing data as reference. The performance is assessed by means of Pearson correlation and root-mean-square error (RMSE) values between the cell type proportions computed by the different deconvolution methods (P_C_; computed proportions; Figure 1) and known compositions (P_E_; expected proportions) of a thousand pseudo-bulk mixtures from each of four different single cell RNA-sequencing datasets (three from human pancreas and one from peripheral blood mononuclear cells (PBMCs)). Furthermore, to evaluate the robustness of our conclusions, different number of cells (cell pool sizes) were used to build the pseudo-bulk mixtures.

**Figure 1.**
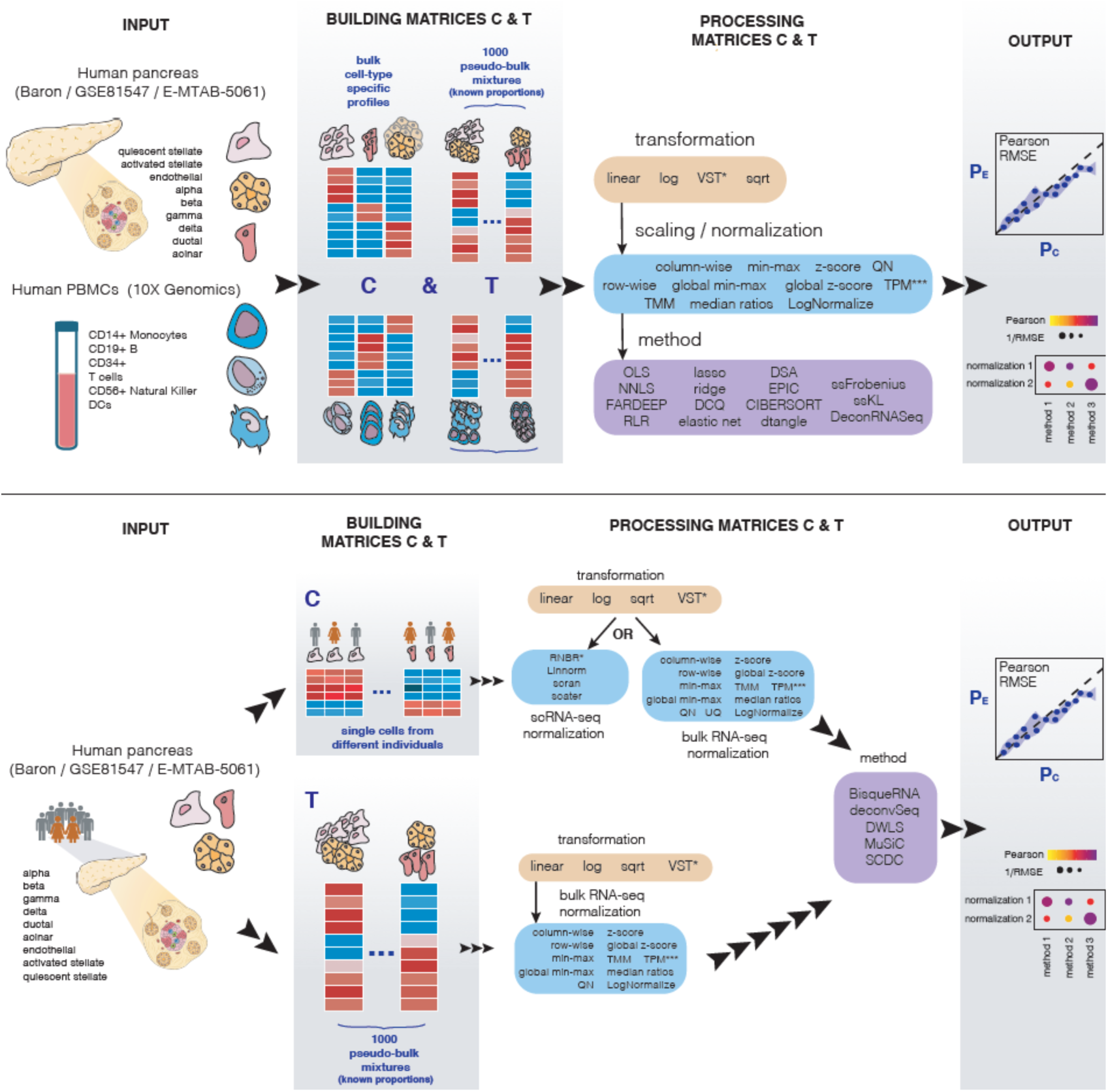
Schematic representation of the benchmarking study. Top panel: workflow for bulk deconvolution methods. Bottom panel: workflow for single-cell methods. In both cases the deconvolution performance is assessed by means of Pearson correlation and root-mean-square error (RMSE). PBMCs = peripheral blood mononuclear cells; log = logarithmic; sqrt = square-root; VST = Variance stabilization transformation. P_E_ = Expected proportions; P_c_ = Computed proportions.

## Results

### Different normalization and methodology combinations have different memory requirements and time consumption

Even though computational resources keep on growing exponentially, memory requirements and time consumption can become important bottlenecks for non-experienced users that may be constrained to limited resources on a personal laptop or for implementations in clinical settings where short processing times are required. While simple logarithmic (log) and square-root (sqrt) data transformations were performed almost instantaneously in R (between 1 and 5 seconds; see Table 1 for information about the number of cells subject to transformation in each single-cell RNA-seq dataset), the variance stabilization transformation (VST) performed using DESeq2^13^ applied to the single-cell RNA-sequencing datasets had high memory requirements and took several minutes to complete (time increasing linearly with respect to the number of cells) (Supplementary Figure 1). Importantly, we used the developer version of DESeq2 v1.25.9, which reduced the running time from quadratic (Suppl. Fig 27 from Soneson *et al*.^14^) to linear with respect of the number of cells.

**Table 1.**
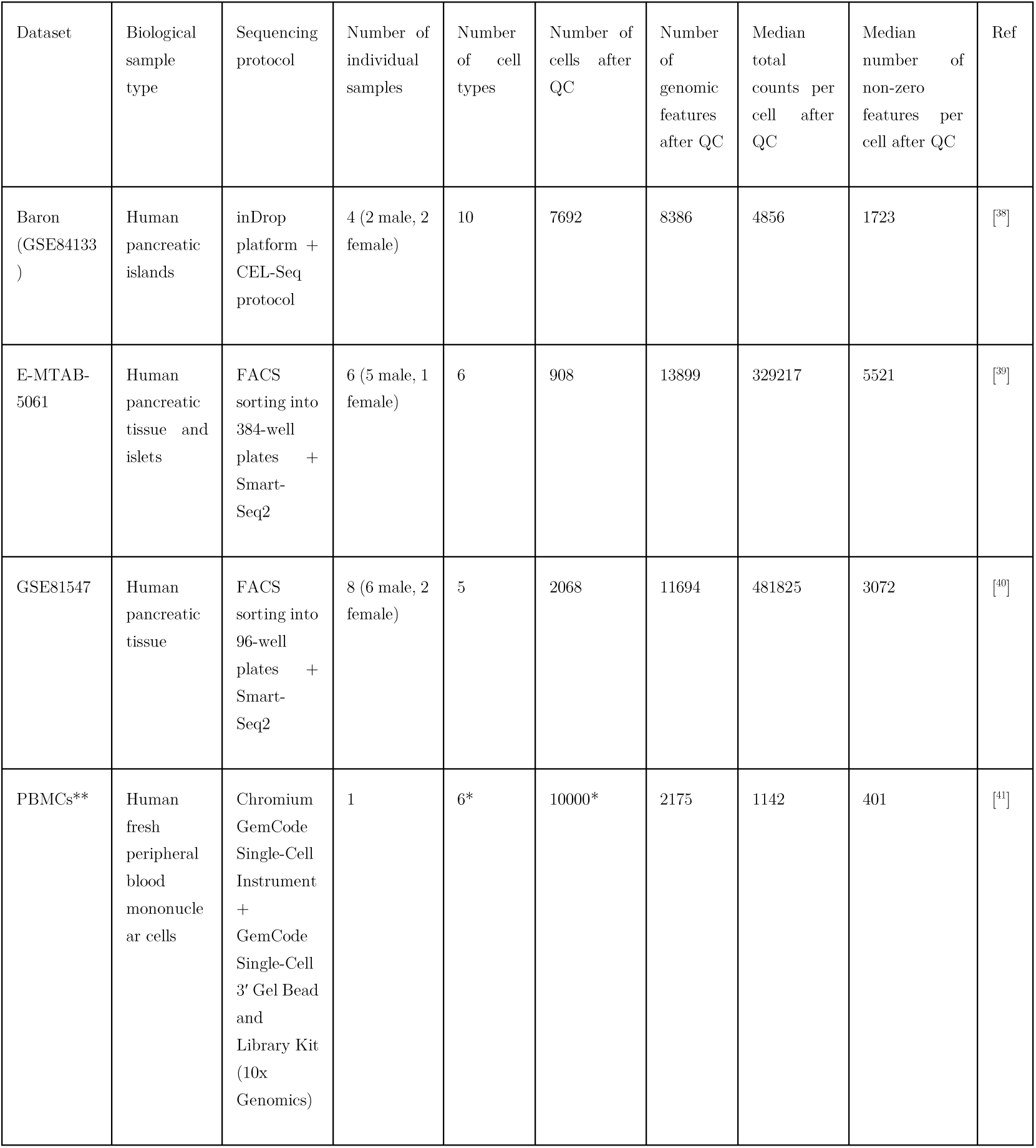
Details of the four datasets used. (*) Since this dataset originally contained six closely related T-cell subtypes (and other people have failed in their attempts of distinguishing them36,37) we re-labelled all cells from these sub-types as “T cells”. Moreover, to reduce the memory and time requirements needed to run all combinations of data transformation, normalization and methodology, we randomly selected 10,000 cells out of the original 68,000. (**) 10X genomics data is not in a public repository but available at: https://support.10xgenomics.com/single-cell-gene-expression/datasets/1.1.0/fresh_68k_pbmc_donor_a

| Dataset | Biological sample type | Sequencing protocol | Number of individual samples | Number of cell types | Number of cells after QC | Number of genomic features after QC | Median total counts per cell after QC | Median number of non-zero features per cell after QC | Ref |
| --- | --- | --- | --- | --- | --- | --- | --- | --- | --- |
| Baron (GSE84133) | Human pancreatic islands | inDrop platform + CEL-Seq protocol | 4 (2 male, 2 female) | 10 | 7692 | 8386 | 4856 | 1723 | [<sup>38</sup>] |
| E-MTAB-5061 | Human pancreatic tissue and islets | FACS sorting into 384-well plates + Smart-Seq2 | 6 (5 male, 1 female) | 6 | 908 | 13899 | 329217 | 5521 | [<sup>39</sup>] |
| GSE81547 | Human pancreatic tissue | FACS sorting into 96-well plates + Smart-Seq2 | 8 (6 male, 2 female) | 5 | 2068 | 11694 | 481825 | 3072 | [<sup>40</sup>] |
| PBMCs** | Human fresh peripheral blood mononuclear cells | Chromium GemCode Single-Cell Instrument + GemCode Single-Cell 3' Gel Bead and Library Kit (10x Genomics) | 1 | 6* | 10000* | 2175 | 1142 | 401 | [<sup>41</sup>] |

We further evaluated the impact of different scaling and normalization strategies as well as the choice of deconvolution method. Although the different scaling/normalization strategies consistently have similar memory requirements, RNBR^15^ and scran^16^ (two single-cell RNA-sequencing specific normalization methods) required up to seven minutes to complete, a 14 fold difference with the other methods, which finished under 30s (Supplementary Figure 2).

The bulk deconvolution methods DSA^17^, ssFrobenius and ssKL^18^ (all implemented as part of the CellMix^19^ R package) had the highest RAM memory requirements, followed by DeconRNASeq^20^. Not surprisingly, the ordinary least squares (OLS^21^) and non-negative least squares (nnls^22^) were the fastest, as they have the simplest optimization problem to solve. For single-cell methods, Dampened Weighted Least Squares (DWLS^23^), which includes an internal marker selection step, resulted in the longest time consumption (6 to 12 hours to complete) whereas MuSiC^24^ and SCDC^25^ finished in 5 to 10 minutes.

### Data transformation has a dramatic impact on the deconvolution results

We investigated the overall performance of each individual deconvolution method across four different data transformations and all normalization strategies (Figure 2; Supplementary Figures 3-4). Maintaining the data in linear scale (“none” transformation, in grey) consistently showed the best results (lowest RMSE values) whereas the logarithmic (in orange) and VST (in green; which also performs an internal complex logarithmic transformation) scale led to a poorer performance, with two to four-fold higher median RMSE values. For a detailed explanation concerning several bulk and single-cell deconvolution methods that could only be applied with a specific data transformation or dataset, please see Supplementary Methods.

**Figure 2.**
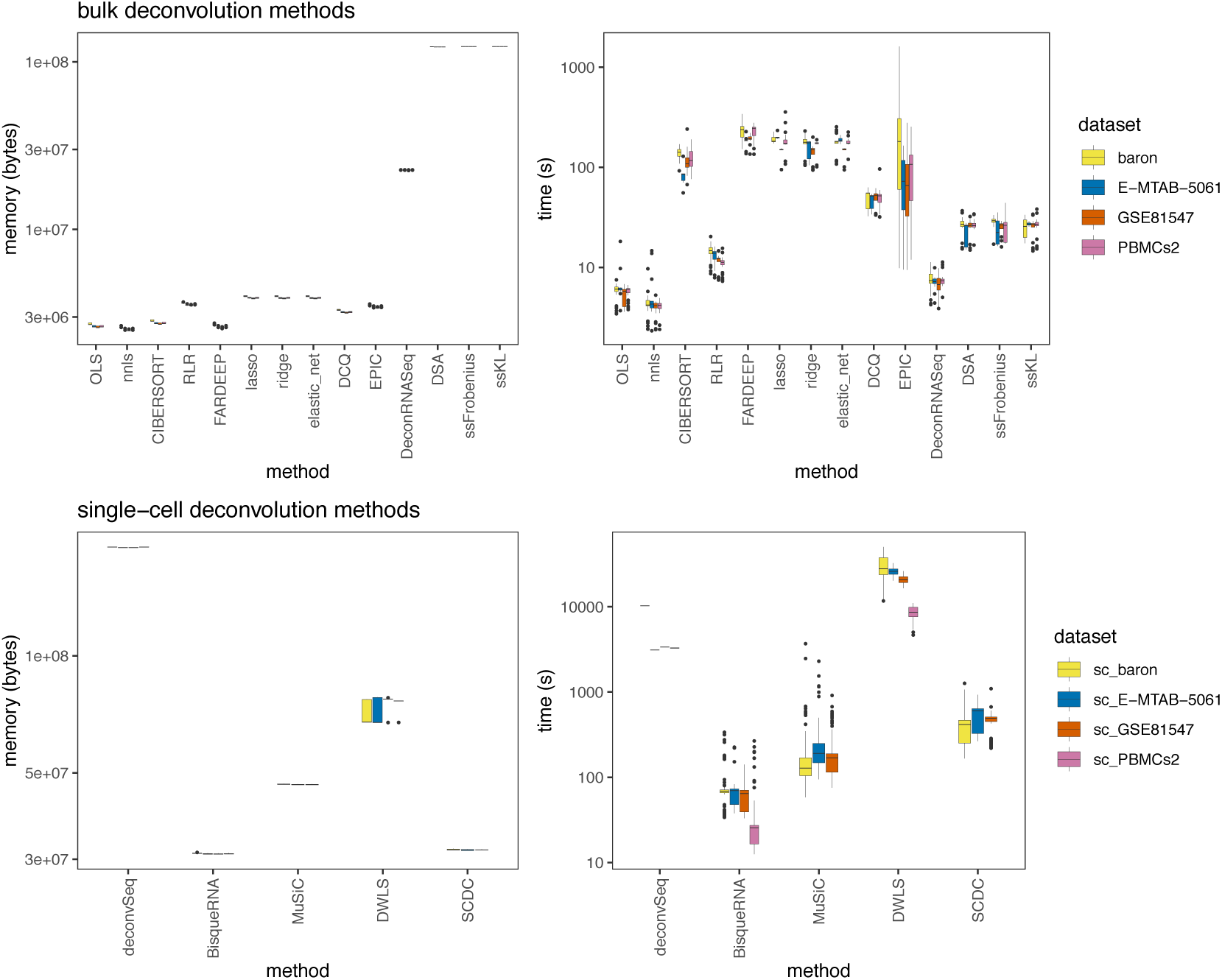
RAM memory (bytes) and time (seconds) requirements for the different bulk (top panel) and single-cell (bottom panel) deconvolution methodologies across datasets with expression values in linear scale (boxplots depict all scaling/normalization strategies across all pseudo-bulk cell pool sizes).

With the exception of EPIC^26^, DeconRNASeq^20^ and DSA^17^, the choice of normalization strategy does not have a substantial impact on the deconvolution results (evidenced by narrow boxplots). These conclusions also hold when repeating the analysis with different pseudo-bulk pool sizes in all datasets tested (collapsing all scaling/normalization strategies and all bulk (Supplementary Figure 5) or single-cell (Supplementary Figure 6) deconvolution methods together). For these reasons, all downstream analyses were performed on datasets in linear scale. Interestingly, there are five bulk (OLS, nnls, RLR, FARDEEP and CIBERSORT) and three single-cell deconvolution methods (DWLS, MuSiC, SCDC) able to achieve very accurate cell type proportions, with median RMSE values lower than 0.05.

### Different combinations of normalization and deconvolution methodologies reveal important differences in performance

From Figure 3 it is clear that different combinations of normalizations and methodologies lead to substantial differences in performance. Focusing on the data in linear scale, Figure 4 delves into the specific method and normalization combinations evaluated in this manuscript. Among the bulk deconvolution methods, least-squares (OLS, nnls), support-vector (CIBERSORT) and robust regression approaches (RLR/FARDEEP) gave the best results across different datasets and pseudo-bulk cell pool sizes (median RMSE values < 0.05; Figure 4a, Supplementary Fig 7). Regarding the choice of normalization/scaling strategy, column min-max and column z-score consistently led to the worst performance. In all other situations, the choice of normalization/scaling strategy had a minimal impact on the deconvolution results for these methods. Of note, quantile normalization always resulted in sub-optimal results in any of the tested bulk deconvolution methods (Figure 4b).

**Figure 3.**
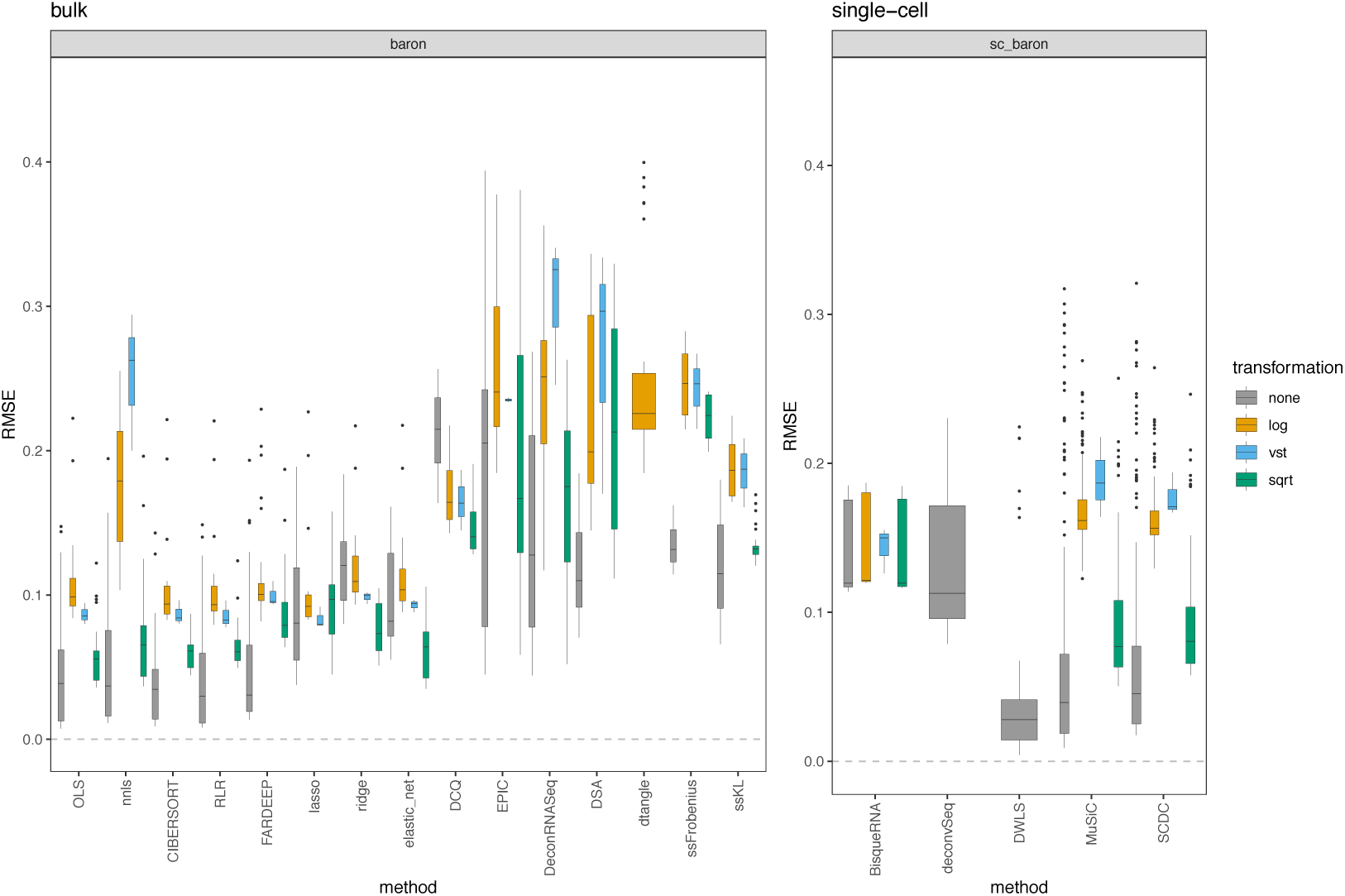
RMSE values between the known proportions in 1000 pseudo-bulk tissue mixtures from the baron dataset (pool size = 100 cells per mixture) and the predicted proportions from the different bulk (left) and single-cell (right) deconvolution methods. Each boxplot contains all normalization strategies that were tested in combination with a given method.

**Figure 4.**
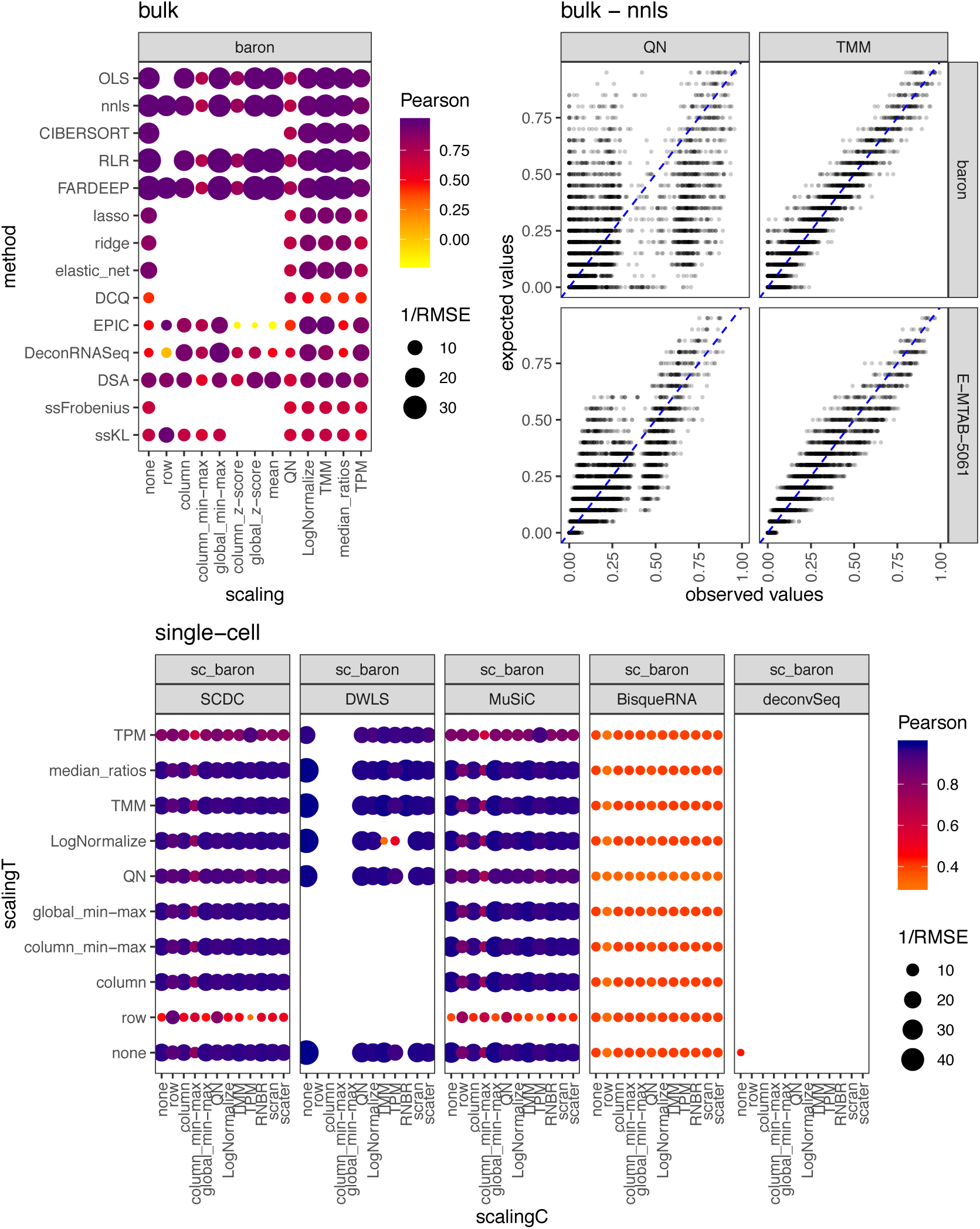
Pearson correlation values between the expected (known) proportions in 1000 pseudo-bulk tissue mixtures in linear scale (pool size = 100 cells per mixture) and the output proportions from the different bulk (a) and single-cell (c) deconvolution methods. The darker the blue and the higher the area of the circle represents higher Pearson and lower RMSE values, respectively. b) Scatter plot showing the impact of the normalization strategy (TMM versus quantile normalization (QN)) comparing the expected proportions (y-axis) and the results obtained through computational deconvolution using nnls (x-axis) for baron and E-MTAB-5061 datasets. Empty locations represent combinations that were not feasible (see Supplementary methods).

Penalized regression approaches including lasso, ridge, elastic net regression and DCQ performed slightly worse than the ones described above (median RMSE ∼ 0.1). As stated in its original publication, EPIC assumes transcripts per million (TPM) normalized expression values as input. We indeed observed that the choice of scaling/normalization has a big impact on the performance of EPIC, with TPM giving the best results.

Quadratic programming (DeconRNASeq), Digital Sorting Algorithm (DSA) and the semi-supervised approaches ssKL and ssFrobenius (using only sets of marker genes, in contrast to the supervised counterparts which use a reference matrix with expression values for the markers) showed the poorest performances with the highest root-mean-square errors and lower Pearson correlation values.

For single-cell deconvolution methods (Figure 4c), we evaluated the different combinations of normalization strategies of both the pseudo-bulk mixtures (“scalingT”, y-axis) and the single-cell expression matrices (“scalingC”, x-axis). DWLS, MuSiC and SCDC consistently showed the highest performance (comparable to the top-performers from the bulk methods, see also Figure 3) across the different choices of normalization strategy (with the exception of row-normalization, column min-max and TPM). While these results are consistent for deconvSeq, MuSiC, DWLS and SDCD regardless of the dataset and pseudo-bulk cell pool size, we observed a substantial performance improvement in BisqueRNA when the pool size increased or when the dataset contained single-cell RNA-sequencing from more individuals (E-MTAB-5061 and GSE81547, with n=6 and 8 respectively) (Supplementary Figure 8). Note that it was not feasible to evaluate all combinations (empty locations in the grid), see Supplementary methods for a detailed explanation.

### The set of markers used in bulk deconvolution methods impacts deconvolution results

Based on the previous results, we wanted to evaluate whether different marker selection strategies had an impact on the deconvolution results starting from bulk expression data in linear scale. To that end we assessed the impact of eight different marker selection strategies (see Methods) on the deconvolution results using bulk deconvolution methods (Figure 5, Supplementary Figure 9). This analysis was not done with the single-cell methods because they do not require marker genes to be known prior to performing the deconvolution.

**Figure 5.**
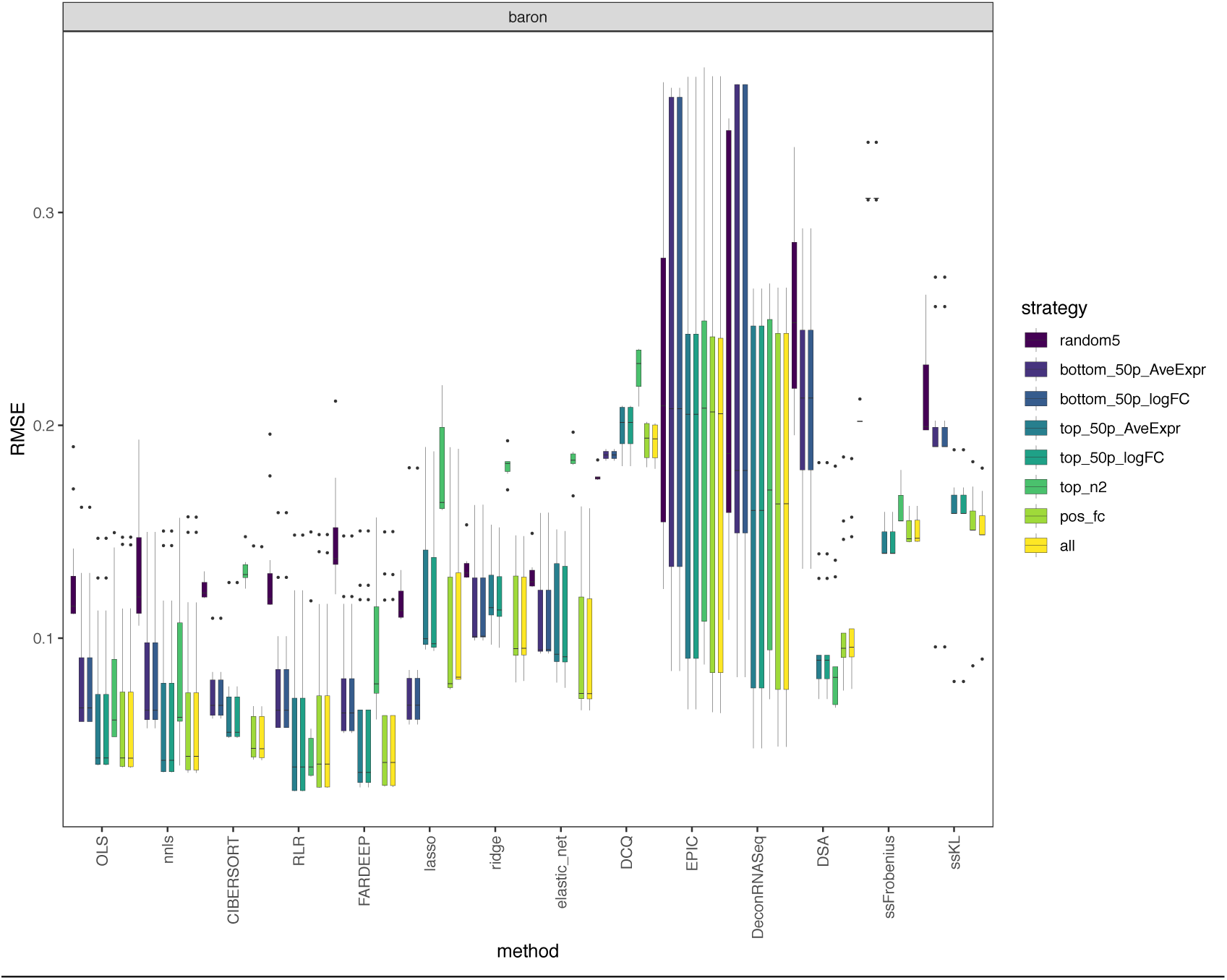
RMSE values between the expected (known) proportions in 1000 pseudo-bulk tissue mixtures (linear scale; pool size = 100 cells per mixture) and the output proportions from the baron dataset, using eight different marker selection strategies. Each boxplot contains all normalization strategies that were tested in combination with a given marker strategy across the different bulk deconvolution methods.

The use of all possible markers (“all” strategy) showed the best performance overall, followed by positive fold-change markers (“pos_fc”; negative fold-change markers are those with small expression values in the cell type of interest and high values in all the others) or those on the top 50% of average expression values (“top_50p_AveExpr”) or log fold-changes (“top_50p_logFC”). As expected, the use of random sets of 5 markers per cell type (“random5”; negative control in our setting) was consistently the worst choice across all datasets regardless of the deconvolution method. Using the bottom 50% of the markers per cell type based on average expression levels (“bottom_50p_AveExpr”) or log fold changes (“bottom_50p_logFC) also led to sub-optimal results. Specifically in the baron and PBMC datasets, the use of the top 2 markers per cell type (“top_n2”) led to a) optimal results when used with DSA; b) similar results as using the bottom_50p_AveExpr or bottom_50p_logFC with ordinary linear regression strategies; c) worse results than random when used with penalized regression strategies (lasso, ridge, elastic_net, DCQ) and CIBERSORT.

### Removing cell types from the reference matrix results in substantially worse deconvolution results compared to reference matrices composed of all cell types present in the mixtures

Based on the results from all the analyses thus far, we decided to evaluate the impact of removing cell types with the data in linear scale and using all available markers (“all” marker selection strategy). Furthermore, we selected nnls and CIBERSORT as representative top-performing bulk deconvolution methods and DWLS and MuSiC as top-performing single-cell methods. To also be able to evaluate the impact of the normalization strategy, we included a representative sample of normalization strategies that result in small RMSE and high Pearson correlation values (see Figure 4 and Supplementary Figures 7-8): column, median ratios, none, TMM and TPM for nnls and CIBERSORT; column, scater, scran, none, TMM and TPM for DWLS and MuSiC.

We assessed the impact of removing a specific cell type by comparing the absolute RMSE values between the ideal scenario where the reference matrix contains all the cell types present in the pseudo-bulk mixtures (leftmost column in Figures 6c-d and 7c-d, with grey label “none”) and the RMSE values obtained after removing one cell type at a time from the reference (all other grey labels).

**Figure 6.**
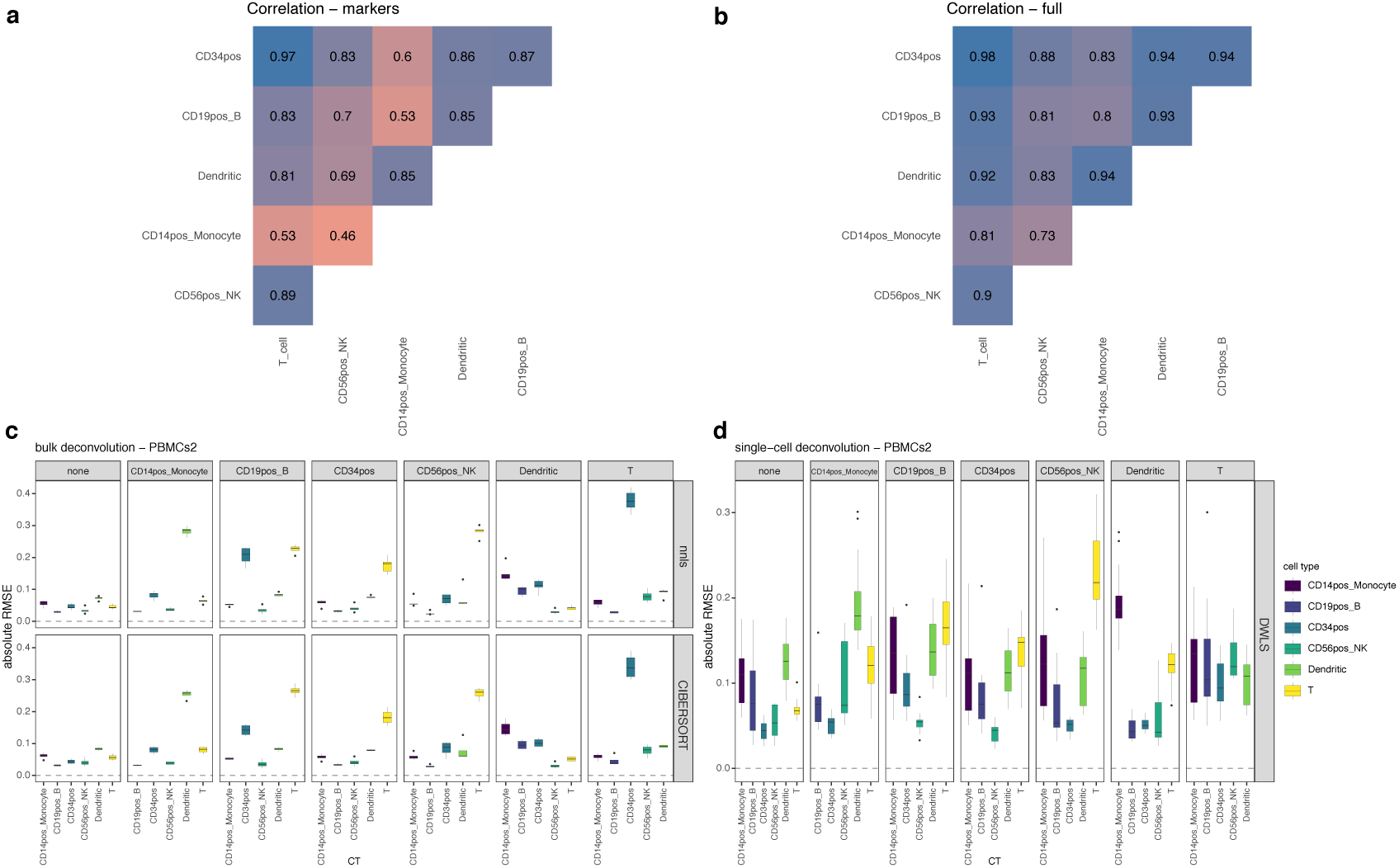
Effect of cell type removal on the deconvolution results using the PBMCs dataset [100-cell pseudo-bulk mixtures in linear scale]. a) pairwise Pearson correlation values between expression profiles for the different cell types, using a subset of the reference matrix containing only the markers used in the bulk deconvolution; b) pairwise Pearson correlation values between complete expression profiles for the different cell types; c) results using bulk deconvolution methods (nnls and CIBERSORT); d) results using single-cell deconvolution methods (only DWLS because the scRNA-seq data comes from only one individual). In c) and d), each grey column represents a specific cell type removed. Each data point conforming a boxplot represents a different scaling/normalization strategy used.

**Figure 7.**
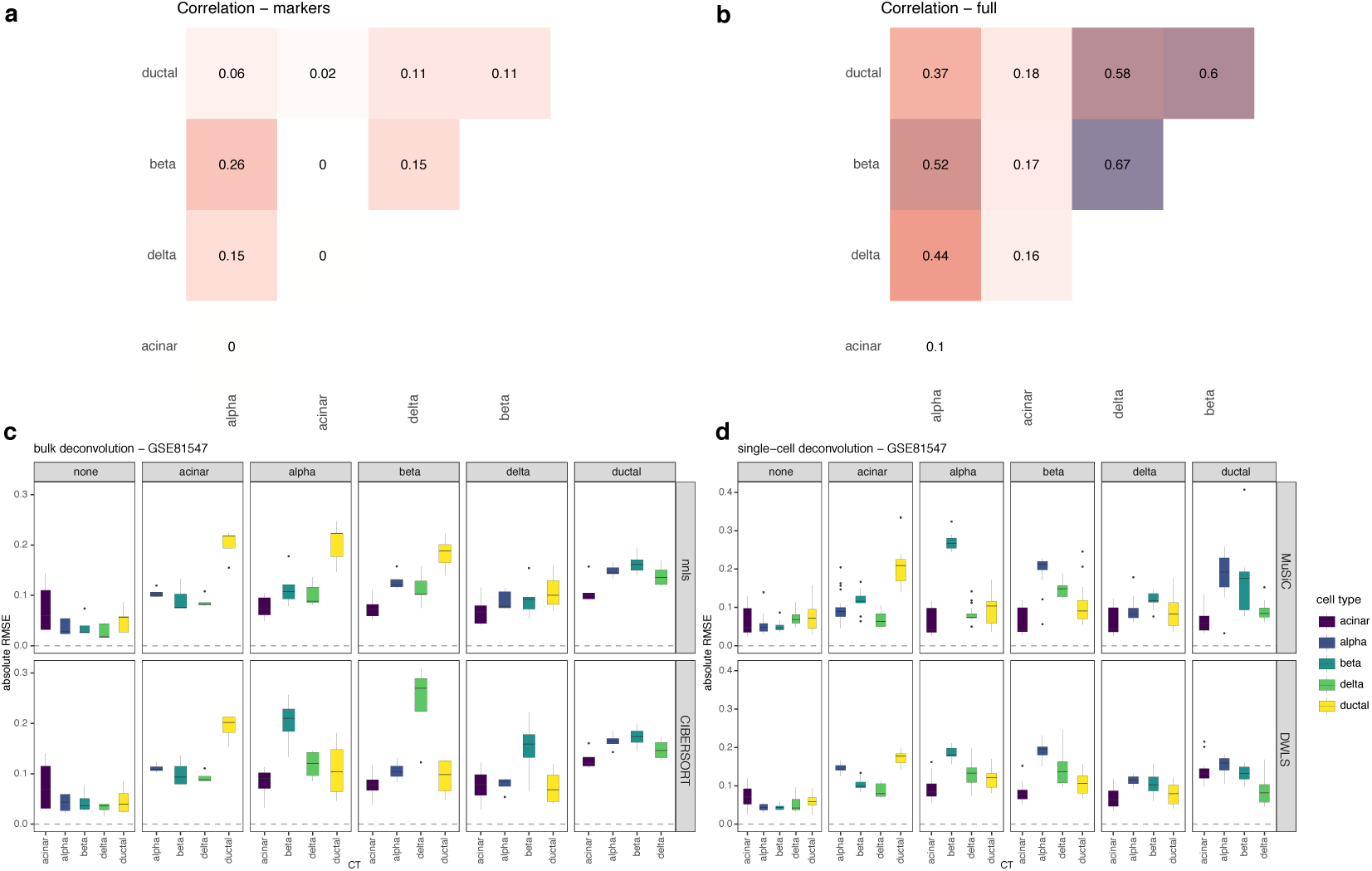
Effect of cell type removal on the deconvolution results using the GSE81547 dataset [100-cell pseudo-bulk mixtures in linear scale]. a) pairwise Pearson correlation values between expression profiles for the different cell types, using a subset of the reference matrix containing only the markers used in the bulk deconvolution; b) pairwise Pearson correlation values between complete expression profiles for the different cell types; c) results using bulk deconvolution methods (nnls and CIBERSORT); d) results using single-cell deconvolution methods (MuSiC and DWLS). In c) and d), each grey column represents a specific cell type removed. Each data point conforming a boxplot represents a different scaling/normalization strategy used.

We then focussed on those cases where the median absolute RMSE values between the results using the complete reference matrix (depicted as “none” in Figures 6c-d and 7c-d) and all other scenarios where a cell type was removed, increased at least 2-fold. In the PBMC dataset (Fig 6c-d), removing CD19+, CD34+, CD14+ or NK cells had an impact on the computed T-cell proportions (between a three and six-fold increase in the median absolute RMSE values, both in bulk and single-cell deconvolution methods). The GSE81547 dataset (Figure 7c-d) shows that removing acinar cells has a dramatic impact in all other cell type proportions. Supplementary Figures 10 and 11 showed the results for baron and E-MTAB-5061 datasets, respectively. Remarkably, no method and normalization combination was able to provide accurate cell type proportion estimates when the reference missed a cell type.

To investigate whether the proportion of the omitted cell type was re-distributed equally among all remaining cell types or only among those that are transcriptionally most similar, we computed pairwise Pearson correlation values between the expression profiles of the different cell types (Figure 6a-b and Figure 7a-b). Figure 6a-b shows that CD14+ monocytes were mostly correlated with dendritic cells (Pearson = 0.85 when computing pairwise correlations on the reference matrix containing only marker genes and 0.94 when using the complete expression profiles from all cell types, respectively) and Figure 6c-d shows that, when removing CD14+ monocytes, the highest RMSE value was found in dendritic cells. Figure 7a-b shows that acinar cells are not correlated with any other cell type (Pearson values close to zero with all other cell types) and Figure 7c-d shows that, when removing acinar cells, all cell type proportions estimates have higher RMSE values compared to the case where no cell type is missing (“none”, leftmost panel). For the baron dataset (Supplementary Figure 10): the removal of ductal cells (highest correlation with quiescent stellate and endothelial cells) led to highest RMSE values for both quiescent stellate and endothelial cells while the removal of endothelial cells (mostly correlated with quiescent stellate, beta and ductal cells) led to the highest RMSE values for quiescent, ductal and beta cells. For the E-MTAB-5061 dataset (Supplementary Figure 11): no cell type is correlated to one another and removing any cell type from the reference matrix led to distorted proportions for all other cell types.

## Discussion

Using both Pearson correlation and RMSE values as measures of the deconvolution performance, we comprehensively evaluated the combined impact of four data transformations, twenty scaling/normalization strategies, seven marker selection approaches and twenty different deconvolution methodologies on four different single-cell RNA-seq datasets. These datasets encompass two different biological sample types (human pancreas and peripheral blood mononuclear cells) and three different sequencing protocols (CEL-Seq, Smart-Seq 2 and GemCode Single-Cell 3′). Additionally, we assessed the impact of using different number of cells when making the pseudo-bulk mixtures and the impact of removing cell types from the reference matrix that were actually present in the mixtures.

Even though the four datasets used throughout this manuscript encompass different sequencing protocols that led to hundred-fold differences in the number of reads sequenced per cell (Table 1), our findings were consistent regardless of the dataset being evaluated or the number of cells used to make the pseudo-bulk mixtures.

The logarithmic transformation is routinely included as a part of the pre-processing of omics data in the context of differential gene expression analysis ^27,28^, but Zhong and Liu^6^ showed that it led to worse results than performing computational deconvolution in the linear (un-transformed) scale. Silverman *et al*.^29^ showed that using log counts per million with sparse data strongly distorts the difference between zero and non-zero values and Townes *et al*.^30^ showed the same when log-normalizing UMIs. Tsoucas *et al*.^23^ showed that when the data was kept in the linear scale, all combinations of three deconvolution methods (DWLS, QP or SVR) and three normalization approaches (LogNormalize from Seurat, Scran or SCnorm) led to a good performance, which was not the case when the data was log-transformed. Here, we assessed the impact of the log transformation on both full-length and tag-based scRNA-seq quantification methods and confirmed that the computational deconvolution should be performed on linear scale to achieve the best performance.

Data scaling or normalization is a key pre-processing step when analysing gene expression data. Data scaling approaches transform the data into bounded intervals such as [0, 1] or [-1, +1]. While being relatively easy and fast to compute, scaling is sensitive to extreme values. Therefore, other strategies that aim to change the observations so that they follow a normal distribution (= normalization) may be preferred. Importantly, these normalizations typically do not result in bounded intervals. In the context of transcriptomics, normalization is needed to only keep true differences in expression. Normalizations such as TPM aim at removing differences in sequencing depth among the samples. We refer the reader to Evans *et al*.^31^, for an in-depth analysis of RNA-seq normalization methods. Vallania *et al*. ^11^ assessed the impact of standardizing (= substracting the mean and dividing by the standard deviation) both the bulk and reference expression profiles into z-scores prior to deconvolution, which is performed by CIBERSORT but not in other methods. They observed high pairwise correlations between the estimated cell type proportions with and without standardizing the data, suggesting a neglectable effect. However, a high Pearson correlation value is not always synonym of a good performance. As already pointed out by Hao *et al*.^32^, high Pearson correlation values can arise when the proportion estimations are accurate (low RMSE values) but also when the proportions differ substantially (high RMSE values), making the correlation metric alone not sufficient to assess the deconvolution performance. Both for bulk and single-cell deconvolution methods, our analyses show that the normalization strategy had little impact (except for EPIC, DeconRNASeq and DSA bulk methods). Of note, quantile normalization (QN), an approach used by default in several deconvolution methods (e.g. FARDEEP, CIBERSORT), consistently showed sub-optimal performance regardless of the method.

Schelker *et al*.^33^ and Racle *et al*.^26^ showed that the origin of the expression profiles had also a dramatic impact on the results, revealing the need of using appropriate cell types coming from niches similar to the bulk being investigated.

Hunt *et al.*^34^ showed that a good deconvolution performance was achieved if the markers being used were predominantly expressed in only one cell type and with the expression in other cell types being in the bottom 25%. Monaco *et al.*^35^ showed similar conclusions when the reference matrix was pre-filtered by removing markers with small log fold change between the first and second cell types with highest expression. In our analyses, markers were selected based on the fold change with respect to the cell type with the second highest expression. Therefore, the pre-filtering proposed by Hunt *et al*. and Monaco *et al*. was already implicitly done. Furthermore, when sub-setting the markers based on their average gene expression or fold changes, those in the top fifty percent led to smaller RMSEs compared to those in the bottom fifty percent (Figure 5).

Wang *et al*.^24^ explored the effect of removing one immune cell type at a time from the reference matrix on the estimation accuracy using artificial bulk expression of six pancreatic cell types (alpha, beta, delta, gamma, acinar and ductal) and removing one cell type from the single-cell expression dataset. They observed that, when a cell type was missing in the reference matrix, MuSiC, NNLS and CIBERSORT did not produce accurate proportions for the remaining cell types. Gong and Szustakowski^20^ also investigated this issue by performing a first deconvolution using DeconRNASeq, then removing the least abundant cell population from the reference/basis matrix, and finally repeating the deconvolution with the new matrix. They observed an uneven redistribution of the signal and observed that some initial proportions became smaller. Moreover, Schelker *et al*.^33^ investigated this phenomenon by looking at the correlation coefficient between the results obtained with the complete reference matrix and the results removing one cell type at a time.

We performed similar analyses for four deconvolution methods (two bulk and two single-cell) and eleven normalization strategies (five for bulk, six for single-cell) on three single-cell human pancreas and one PBMC dataset, keeping the data in linear scale. We observed both cases where the choice of normalization strategy had no impact and other cases where it did. Interestingly, the removal of specific cell types did not affect all other cell types equally. Both bulk and single-cell deconvolution methods showed similar trends when removing specific cell types. However, there were some discrepancies in the RMSE values (e.g. removal of beta cells had a substantial impact on the proportions of delta cells but CIBERSORT showed three times higher RMSE values compared to either nnls, MuSiC or DWLS). This may be explained by the fact that for bulk deconvolution methods, we removed both the cell type expression profile and its marker genes from the reference matrix whereas for the single-cell methods, only the cells from the specific cell type were excluded, without applying extra filtering on the genes (MuSiC, SCDC) or because a different signature was internally built (DWLS).

Schelker *et al*. found that B cell and dendritic cell proportions were affected by removing macrophages or monocytes whereas NK cell proportions were affected by removing T cells. Sturm *et al.*, also reported the impact of removing CD8+ T cells on NK cell proportions. Our results on the PBMC dataset agree with those from Schelker *et al*. and Sturm *et al*. but also include novel insights: removing CD19+ B-cells, CD34+, CD14+ monocytes or NK cells had an impact on the computed T-cell proportions and removing CD19+ B-cells, CD56+ NK or T cells had an impact on CD34+ cell proportions.

Furthermore, we found a direct association between the correlation values among the cell types present in the mixtures and the effect of removing a cell type from the reference matrices. Specifically, we hypothesize that: a) removing a cell type that is barely or completely uncorrelated (Pearson < 0.2) to all other cell types remaining in the reference matrix has a dramatic impact in the cell type proportions of all other cell types; b) removing a cell type that was strongly positively correlated (Pearson > 0.6) with one or more cell types still present in the reference matrix leads to distorted estimates for the most correlated cell type(s).

EPIC^26^ shows a first attempt in alleviating this problem by considering an unknown cell type present in the mixture. Nevertheless this is currently restricted to a cancer setting, using markers of non-malignant cells that are not expressed in cancer cells.

## Methods

### Dataset selection and quality control

Four different datasets coming from different single-cell isolation techniques (FACS and droplet-based microfluidics) and encompassing both full-length (Smart-Seq2) and tag-based library preparation protocols (3’-end with UMIs) were used throughout this article (see Table 1). After removing all genes (rows) full of zeroes or with zero variance, those cells (columns) with library size, mitochondrial content or ribosomal content further than three median absolute deviations (MADs) away were discarded. Next, only genes with at least 5% of all cells (regardless of the cell type) with a UMI or read count greater than 1 were kept. Finally, we retained cell types with at least 50 cells passing the quality control step and, by setting a fixed seed and taking into account the number of cells across the different cell types, each dataset was further split into “training” and “testing” datasets with a similar distribution of cells per cell type. Regarding E-MTAB-5061: cells with “not_applicable”, “unclassified” and “co-expression_cell” labels were excluded and only cells coming from six healthy patients (non-diabetic) were kept.

After quality control, we made two-dimensional t-SNE plots for each dataset. When adding coloured labels both by cell type and donor (Suppl. Fig 12), the plots showed consistent clustering by cell type rather than by donor, indicating an absence of batch effects.

### Generation of reference matrices for the deconvolution

Using the “training” splits from the previous section, the mean count across all individual cells from each cell type was computed for each gene, constituting the original (un-transformed and un-normalized) reference matrix (C in equation (I) from section “Computational deconvolution: formulation and methodologies”) and were used as input for the bulk deconvolution methods described in that section. Importantly, the “training” splits without applying the mean collapsing step were used by the single-cell deconvolution methods and for the marker selection step.

### Cell-type specific marker selection

TMM normalization (edgeR package^42^) was applied to the original (linear) scRNA-seq expression datasets and limma-voom^43^ was used to find out marker genes. Only genes with positive count values in at least 30% of the cells of at least one cell type were retained. Among the retained ones, those with absolute fold changes greater or equal to 2 between the first and second cell types with highest expression and BH adj p-value < 0.05 were kept as markers in all three pancreatic datasets. Since the PBMCs contained more closely related cell types, the fold-change threshold was lowered to 1.5.

Once the set of markers was retrieved, the following approaches were evaluated: i) “all”: use of all markers found following the procedure described in the previous paragraph; ii) “pos_fc”: using only markers with positive fold-change (=over-expressed in cell type of interest; negative fold-change markers are those with small expression values in the cell type of interest and high values in all the others); iii) “top_n2”: using the top 2 genes per cell type with the highest log fold-change; iv) “top_50p_logFC”: top 50% of markers (per CT) based on log fold-change; v) “bottom_50p_logFC”: bottom 50% of markers based on log fold-change; vi) “top_50p_AveExpr”: top 50% of markers based on average gene expression (baseline expression); vii) “bottom_50p_AveExpr”: low 50% based on average gene expression; viii) “random5”: for each cell type present in the reference, five genes that passed quality control and filtering were randomly selected as markers.

### Generation of thousands of artificial pseudo-bulk mixtures

Using the “testing” datasets from the quality control step, we generated matrices containing 1,000 pseudo-bulk mixtures (matrix T in equation (I) from **“**Computational deconvolution: formulation and methodologies”) by adding up count values from the randomly selected individual cells. The minimum number of cells used to create the pseudo-bulk mixtures (pool size) was 100 and the maximum was determined by the second most abundant cell type (rounded down to the closest hundred, to avoid non-integer numbers of cells) in each of the four datasets. When the difference between the minimum and maximum values was greater than or equal to 200, three different pool sizes were created by rounding up the mean value between both extremes to the closest hundred (n = 100, 700 and 1200 for Baron; n = 100, 300 and 400 for PBMCs). Due to this constraint, only two pool sizes were feasible for GSE81547 (n = 100 and 200) and one for E-MTAB-5061 (n = 100). Each (feasible) pseudo-bulk mixture was created by randomly selecting the number of cell types to be present (between 3, 4 and 5) and their identities, followed by choosing the cell type proportion assigned to each cell type (enforcing a sum-to-one constraint) among all possible proportions between 0.05 and 1, in increasing intervals of 0.05. Finally, once the amount of cells to be picked up from specific cell types was determined, the cells were randomly selected (without replacement).

### Data transformation and normalization

The next step is applying four different data transformations to: i) the un-transformed and un-normalized reference matrix C; ii) the un-transformed and un-normalized single-cell “training” splits and iii) the un-transformed and un-normalized matrix T containing the 1000 pseudo-bulk mixtures.

Since count data from both bulk and single-cell RNA-seq show the phenomenon of over-dispersion^42,44^, the following data transformations were chosen: a) leave the data in the original (linear) scale; b) use the natural logarithmic transformation (with the log1p function in R^45^); c) use the square-root transformation; d) variance-stabilizing transformation (VST). The second and third are simple and commonly used transformations aiming at reducing the skewness in the data due to the presence of extreme values^28^ and stabilizing the variance of Poisson-distributed counts^46^, respectively. VST (using the varianceStabilizingTransformation function from DESeq2) removes the dependence of the variance on the mean, especially important for low count values, while simultaneously normalizing with respect to library size^13^.

Each transformed output file was further scaled/normalized with the approaches listed on Table 2. The mathematical implementation can be found at the original publications (“Ref” column) and in our GitHub repository (http://github.com/favilaco/deconv_benchmark). Due to the sparsity of the single-cell RNA-seq matrices (most genes with zero counts), the UQ normalization failed (all normalization factors were infinite or NA values) and thus was eventually not included in downstream analyses. TMM includes an additional step that uses the normalization factors to obtain normalized counts per million. LogNormalize and Linnorm include an additional exponentiation scale after normalization in order to transform the output data back into linear scale. Median of ratios can only be applied to integer counts in linear scale.

**Table 2.** Detailed description of different scaling/normalization approaches used in the benchmarking.

| Scaling/normalization method | Single-cell specific | Output containing negative values | Output bounded in [0,1] interval | Reference |
| --- | --- | --- | --- | --- |
| Column-wise (=“Total count” or library size normalization) | no | no | yes | [47] |
| Column min-max | no | no | yes | [48] |
| Column z-score | no | yes | no | [49] |
| Row-wise | no | no | yes | [50] |
| Global min-max | no | no | yes | [48] |
| Global z-score | no | yes | no | [49] |
| Quantile<br>normalization (QN) | no | no | no | [51] |
| Upper quartile (UQ) | no | no | no | [52] |
| Transcripts per<br>million (TPM) | no | no | no | [53] |
| Trimmed mean of M-values (TMM) | no | no | no | [54] |
| LogNormalize | no | no | no | [55] |
| Median of ratios | no | no | no | [13] |
| Scran | yes | no | no | [16] |
| Scater | yes | no | no | [56] |
| Linnorm | yes | no | no | [57] |
| RNBR | yes | no | no | [15] |

### Computational deconvolution: formulation and methodologies

The deconvolution problem can be formulated as T = C · P (I) ^5^, where T = measured expression values from bulk heterogeneous samples; C = cell type-specific expression values and P = cell-type proportions. Specifically, T represents the 1000 pseudo-bulk mixtures from “Generation of thousands of artificial pseudo-bulk mixtures” and C is the reference matrix from “Cell-type specific marker selection and generation of reference matrices for the deconvolution”. In the context of this article, the goal is to obtain P using T and C as input.

Fifteen bulk deconvolution methods a have been evaluated, including two traditional (ordinary least squares (OLS^21^) and non-negative least squares (NNLS^22^)) and one weighted least squares method (EPIC^26^); two robust regression (FARDEEP^58^, RLR^59^), one support-vector regression (CIBERSORT^9^) and four penalized regression (ridge, lasso, elastic net^60^ and Digital Cell Quantifier (DCQ^61^)) approaches; one quadratic programming (DeconRNASeq^20^), one method that models the problem in logarithmic scale (dtangle^34^) and three methods included in the CellMix R package^19^: Digital Sorting Algorithm (DSA^17^) and two semi-supervised non-negative matrix factorization methods (ssKL and ssFrobenius^18^). Furthermore, five single-cell deconvolution methods have been evaluated: deconvSeq^62^, MuSiC^24^, DWLS^23^, Bisque^63^ and SCDC^25^. We refer the reader the original publications and our Github repository (http://github.com/favilaco/deconv_benchmark) for details about their implementation.

### Measures of deconvolution performance

Changes in memory were assessed with the mem_change function from the pryr package^64^ and the elapsed time was measured with the proc.time function (both functions executed in R v.3.6.0).

We computed both the Pearson correlation values and the root-mean-square error (RMSE) between cell type proportions from thousands of pseudo-bulk mixtures with known composition and the output from different deconvolution methods for each combination of data transformation, scaling/normalization choice and deconvolution method. Higher Pearson correlation and low RMSE values correspond to a better deconvolution performance.

### Evaluation of missing cell types in the reference matrix C

For every cell type removed, the deconvolution was applied only to mixtures where the missing cell type was originally present. For bulk deconvolution methods, the marker genes of the cell type that was removed from the reference were also excluded (single-cell methods did not require a priori marker information).

## Conclusion and future perspectives

The three most relevant factors affecting the deconvolution results are: i) the data transformation, ii) all cell types being part of the mixtures must be represented in the reference matrix and, for bulk deconvolution methods, iii) a sensible marker selection strategy.

When performing a deconvolution task, we advise users to: a) keep their input data in linear scale; b) select any of the scaling/normalization approaches described here with exception of row scaling, column min-max, column z-score or quantile normalization; c) choose a regression-based bulk deconvolution method (e.g. nnls, CIBERSORT or FARDEEP) and also perform the same task in parallel with DWLS, MuSiC or SCDC if single-cell data is available; d) use a stringent marker selection strategy that focuses on differences between the first and second cell types with highest expression values; e) use a comprehensive reference matrix that include all relevant cell types present in the mixtures.

Finally, as more scRNA-seq datasets become available in the near future, its aggregation (while carefully removing batch effects) will increase the robustness of the reference matrices being used in the deconvolution and will fuel the development of methodologies similar to SCDC, which allows direct usage of more than one scRNA-seq dataset at a time.

## Acknowledgements

We would like to acknowledge Evan Benn, Derrick Lin, Vikkitharan Gnanasambandapillai and Manuel Sopena Ballesteros for their IT support. This work was supported by the Concerted Research Actions from Ghent University (BOF.DOC.2017.0026.01) and a scholarship for a long stay abroad (V440318N) from the Fund for Scientific Research Flanders (FWO) to FAC.

## Author contributions

FAC conducted all the analyses under the supervision of PM, KDP and JP and FAC wrote the manuscript. JAH contributed to the quality control assessment of the single-cell RNA-seq data. KDP, PM and JAH reviewed and edited the manuscript. All authors read and approved the final manuscript.

## Author information

FAC is a PhD student in the Department of Pediatrics and Medical Genetics, Faculty of Medicine and Health Sciences, Ghent University (Belgium). PM and KDP are Professors in the Department of Pediatrics and Medical Genetics, Faculty of Medicine and Health Sciences, Ghent University (Belgium). JAH is a PhD student at the Institute for Molecular Bioscience, University of Queensland. JP is the head of the Garvan-Weizmann Centre for Cellular Genomics and head of the Computational Genomics Group at the Garvan Institute of Medical Research in Sydney (Australia).

