## Supplementary material for "Comprehensive benchmarking of computational deconvolution of transcriptomics data"

#### Supplementary Methods

Incompatible data transformations or normalizations with several deconvolution methods

Global and column z-score normalizations generated negative values, making it incompatible with the single-cell deconvolution methods and with bulk deconvolution methods such as DeconRNASeq, ssKL, ssFrob and DSA. Quantile normalization is used by default in FARDEEP but we disabled it to observe the impact of other scaling/normalization strategies. Row scaling led to singular matrices (several rows were identical; determinant = 0) and thus methods such as robust linear regression (RLR) failed. Linnorm normalization performs an internal logarithmic transformation step, so it is not compatible with logarithmic, square-root and VST transformed input data. CIBERSORT performs and internal z-score standardization of the input matrices prior to fit the support vector regression. The glmnet function used in penalized regression approaches such as ridge, lasso, elastic net and DCQ, includes an internal standardization step (=predictors to be scaled as z-scores) to ensure that the penalty affects each coefficient equally. DSA, ssFrobenius and ssKL can only be applied to data in linear scale [http://web.cbio.uct.ac.za/~renaud/CRAN/web/CellMix/gedAlgorithm.ssKL.html] whereas dtangle only accepts input matrices in logarithmic scale. ssFrobenius performs an internal mean-centering step of each signature separately whereas in ssKL no re-scaling is performed at all. MuSiC and SCDC could not be tested using PBMCs because n=1 (they are “multi-subject” methods). deconvSeq is formulated as a generalized linear model that accounts for the quadratic relationship between the mean and the variance in RNA-seq count data using the log link function for a negative binomial distribution, so it is not compatible with logarithmic, square-root and vst transformed input data, and it requires the input to be un-normalized. DWLS includes an internal log2 transformation step followed by differential gene expression analysis (internal marker selection step) with Model-based Analysis of Single-cell Transcriptomics (MAST)[1]. For these reasons, only single-cell input data in linear scale and normalization strategies not generating negative or bounded values were compatible with DWLS.

Explicit versus implicit non-negativity and sum-to-one constraints

For some methods, the output needed to be explicitly ("E" in the table below) modified after the deconvolution to enforce only positive proportions (non-negativity constraint: negative proportions were set to 0) and that they sum to one. For others, these constraints were implicitly ("I" in the table below) included and the output was left unchanged.

**Supplementary Table 1** – Explicit (E) versus implicit (I) non-negativity and sum-to-one constraints

| deconvolution method | non-negativity | sum-to-one |
|---|---|---|
| OLS | E | E |
| NNLS | I | E |
| FARDEEP | I | E |
| RLR | E | E |
| lasso | E | E |
| ridge | E | E |
| elastic net | E | E |
| DCQ | E | E |
| DSA | E | E |
| EPIC | I | I |
| dtangle | I | I |
| DeconRNASeq | I | I |
| CIBERSORT | I | I* |
| ssFrobenius | I | I |
| ssKL | I | I |
| deconvSeq | I | I |
| MuSiC | I | I |
| SCDC | I | I |
| Bisque | I | I |
| DWLS | E | E** |

(*) Users can select "absolute" mode to remove the sum-to-one constraint or to use a signature score as output.

(**) It only included the implicit sum-to-one constraint. Thus, to enforce both constraints, we artificially enforced the non-negativity constraint followed by sum-to-one.

### Supplementary Figures

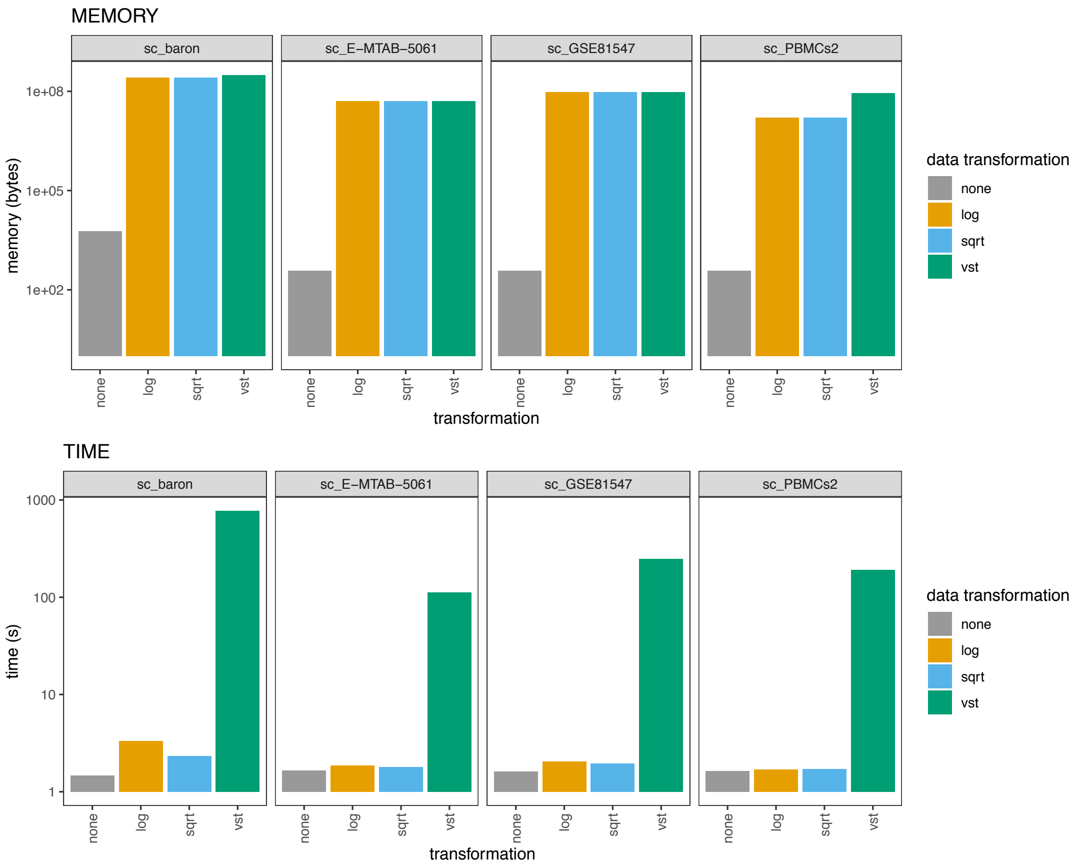


**Supplementary Figure 1** – RAM memory (bytes) and time requirements (seconds) for the different transformations across datasets. "none" represents the data un-transformed, in linear scale; log = logarithmic; sqrt = square-root; vst = variance stabilization transformation.

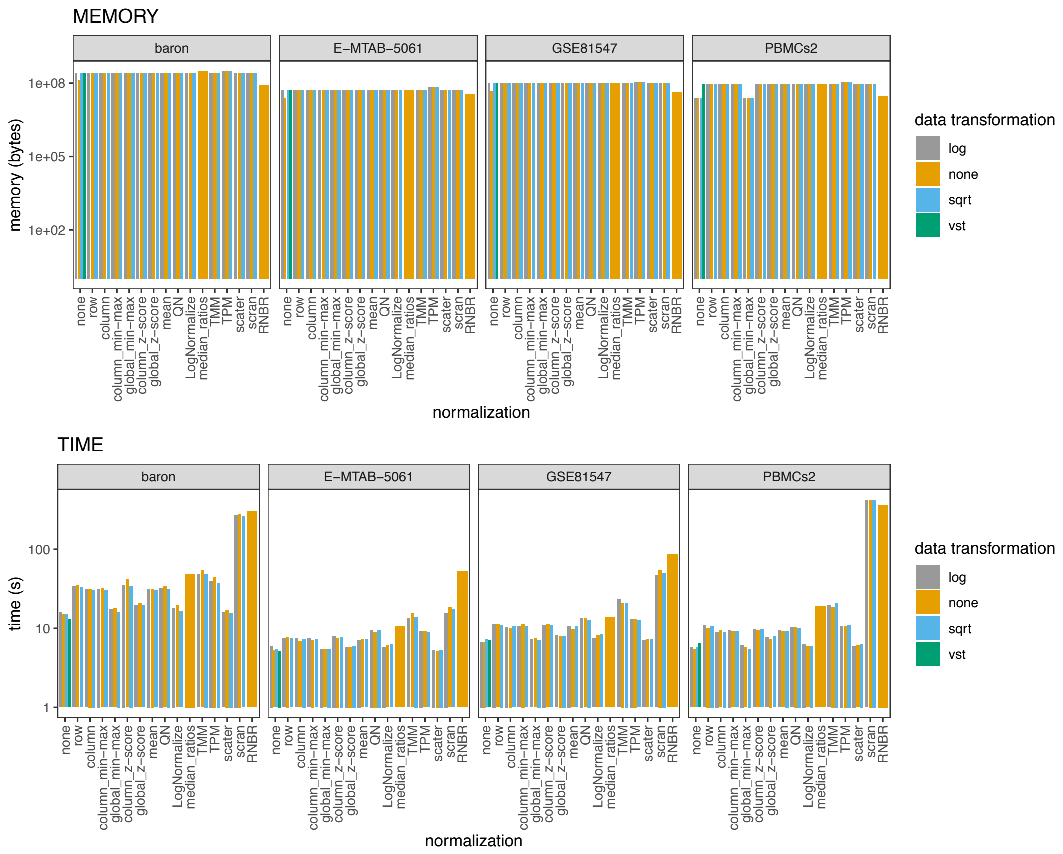


**Supplemental Figure 2** – RAM memory (bytes) and time requirements (seconds) for the different scaling/normalization strategies across different single-cell datasets.

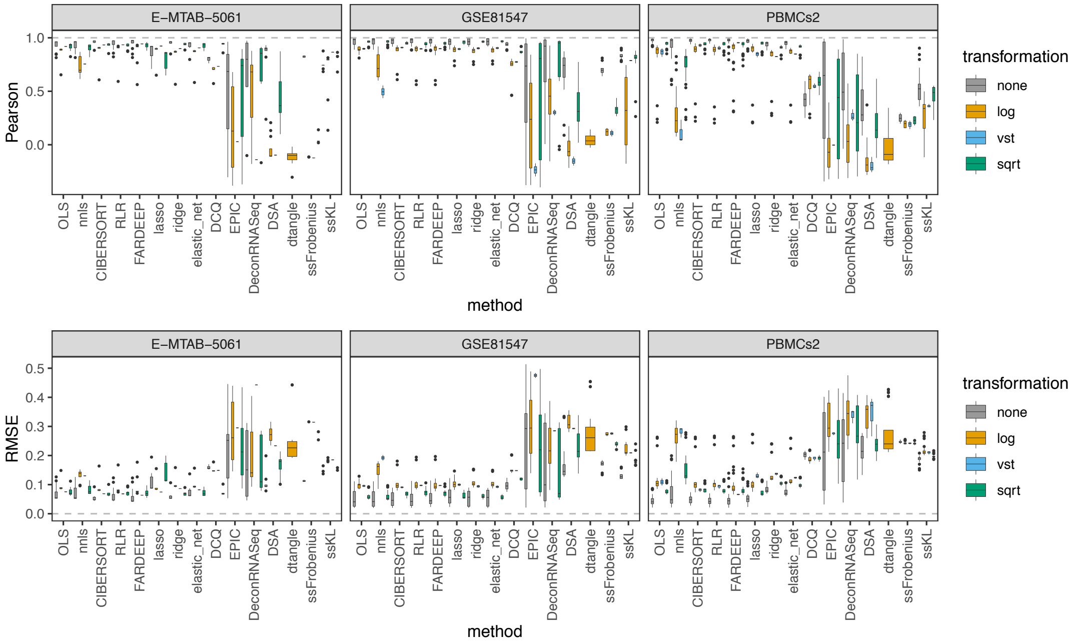


**Supplementary Figure 3** – Pearson correlation (top panel) and RMSE values (bottom panel) between the known proportions in 1000 pseudo-bulk tissue mixtures from the E-MTAB-5061, GSE81547 and PBMCs datasets (pool size = 100 cells per mixture) and the predicted proportions from the different bulk deconvolution methods.

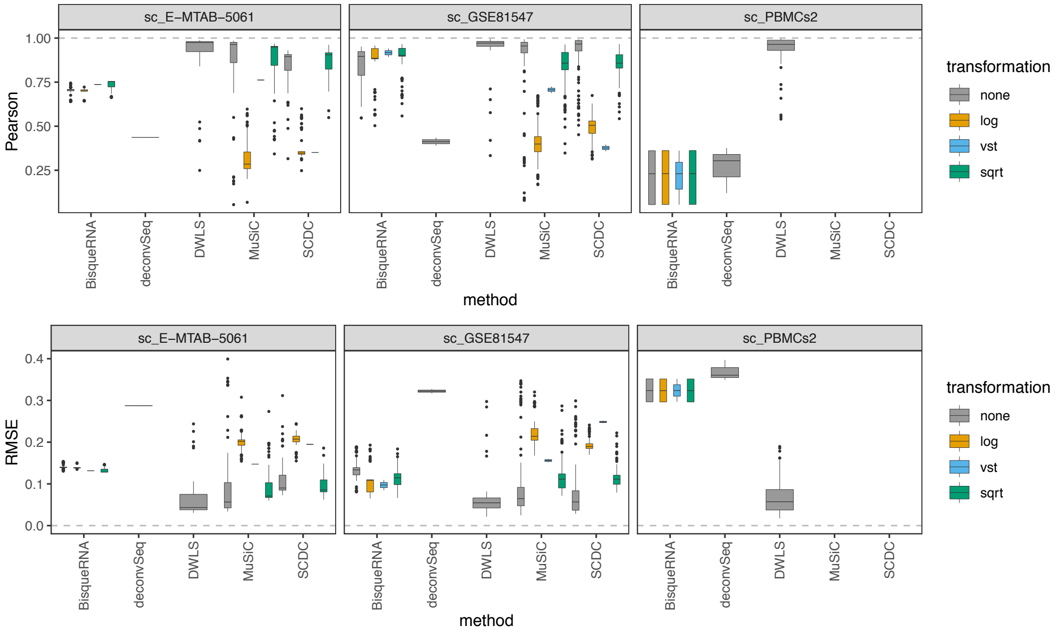


**Supplementary Figure 4** – Pearson correlation (top panel) and RMSE values (bottom panel) between the known proportions in 1000 pseudo-bulk tissue mixtures from the E-MTAB-5061, GSE81547 and PBMCs datasets (pool size = 100 cells per mixture) and the predicted proportions from the different single-cell deconvolution methods. MuSiC and SCDC were not applicable to the PBMC dataset because it requires the number of samples to be greater than one. Each boxplot contains all normalization strategies that were tested in combination with a given method.

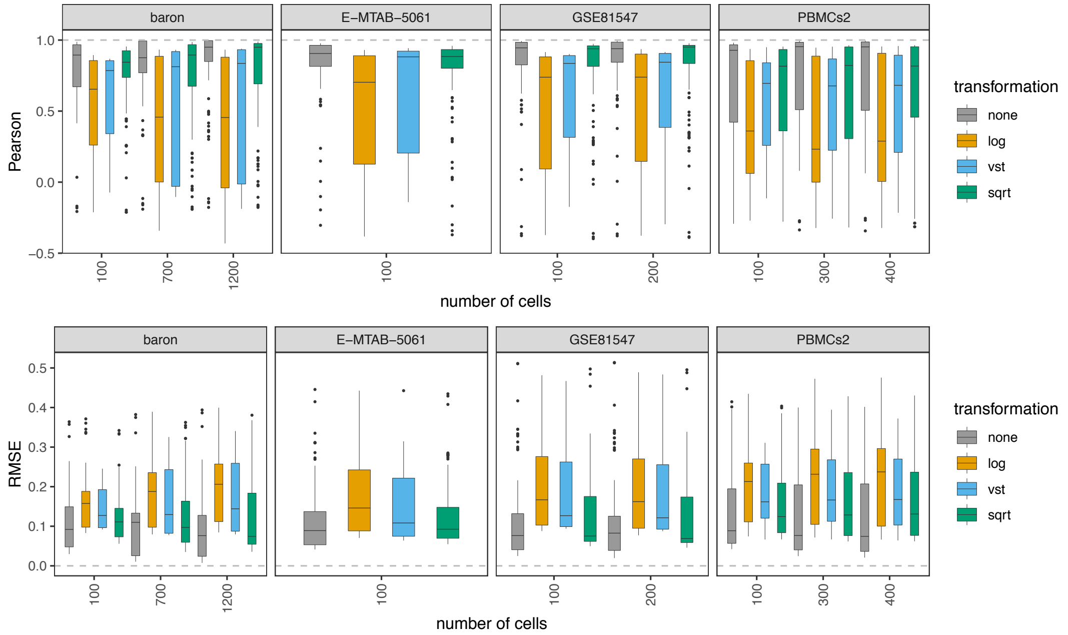


**Supplementary Figure 5** – Pearson correlation (top panel) and RMSE values (bottom panel) between the known proportions in 1000 pseudo-bulk tissue mixtures and the predicted proportions from the different bulk deconvolution methods. Each boxplot contains all combinations of method and normalization strategies that were tested with a given cell pool size.

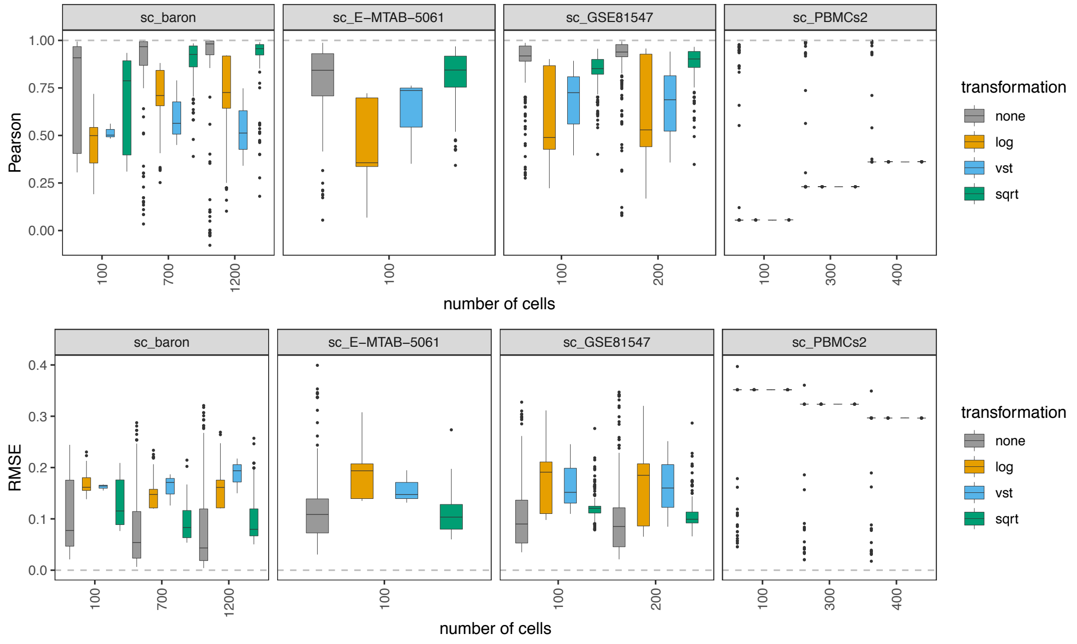


**Supplementary Figure 6** – Pearson correlation (top panel) and RMSE values (bottom panel) between the known proportions in 1000 pseudo-bulk tissue mixtures and the predicted proportions from the different single-cell deconvolution methods. Each boxplot contains all combinations of method and normalization strategies that were tested with a given cell pool size.

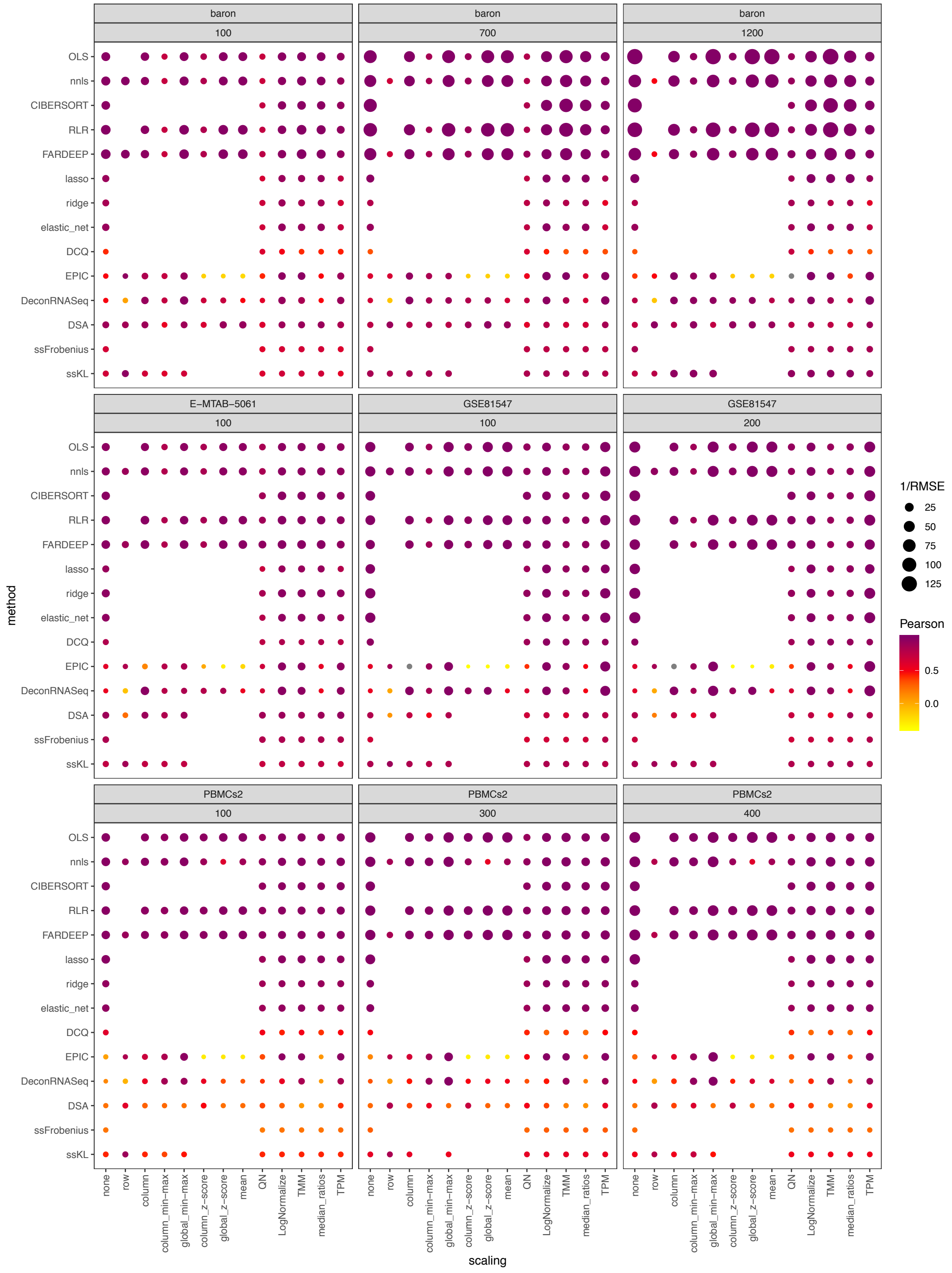


**Supplementary Figure 7** – Pearson correlation values between the expected (known) proportions in 1000 pseudo-bulk tissue mixtures in linear scale (several pool sizes and datasets, as depicted in the grey labels) and the output proportions from the different bulk deconvolution methods. The darker the blue and the higher the area of the circle represents higher Pearson and lower RMSE values, respectively.

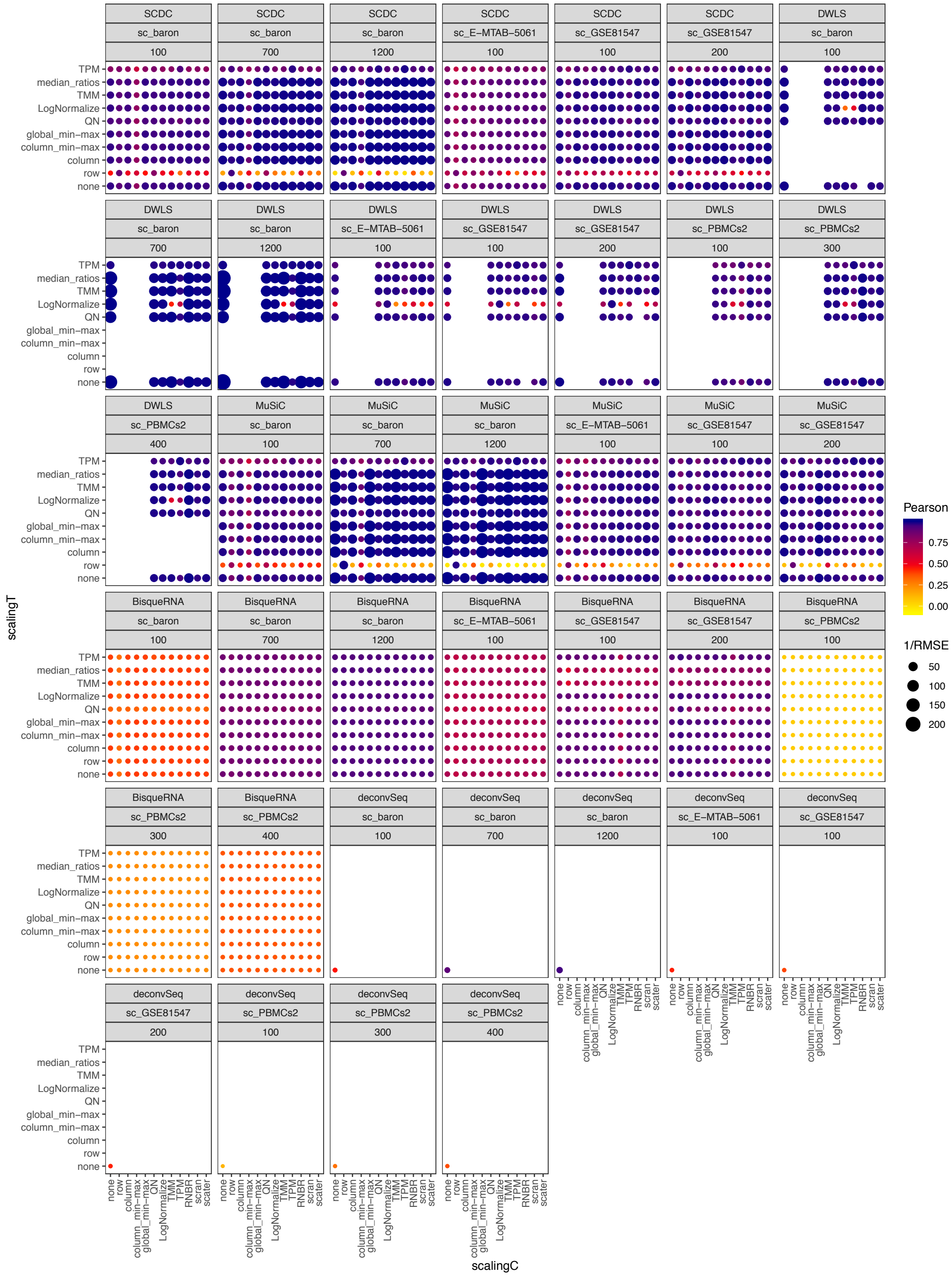


**Supplementary Figure 8** – Pearson correlation values between the expected (known) proportions in 1000 pseudo-bulk tissue mixtures in linear scale (several pool sizes and datasets, as depicted in the grey labels) and the output proportions from the different single-cell deconvolution methods. The darker the blue and the higher the area of the circle represents higher Pearson and lower RMSE values, respectively.

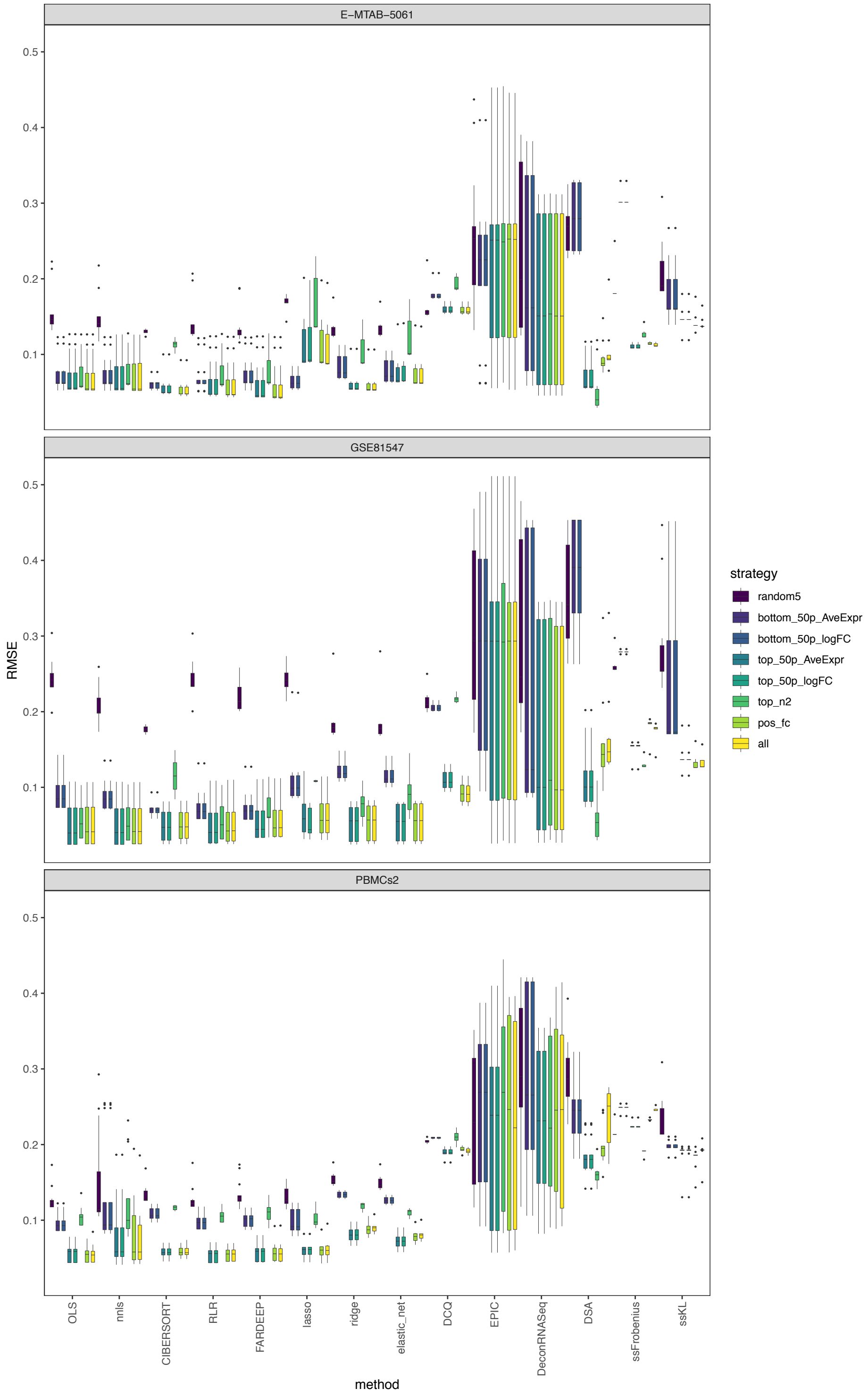


**Supplementary Figure 9** – RMSE values between the expected (known) proportions in 1000 pseudo-bulk tissue mixtures (linear scale; pool size = 100 cells per mixture) and the output proportions from the E-MTAB-5061, GSE81547 and PBMCs datasets, using eight different marker selection strategies. Each boxplot contains all normalization strategies that were tested in combination with a given marker strategy across the different bulk deconvolution methods.

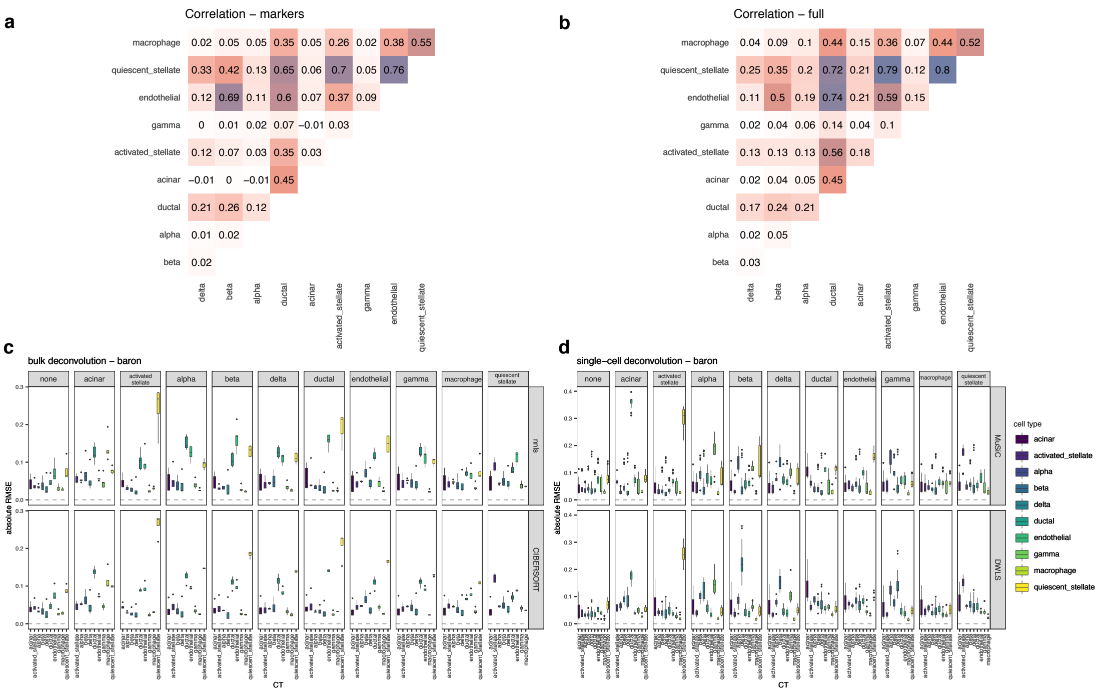


**Supplementary Figure 10** – Effect of cell type removal on the deconvolution results using the baron dataset [100-cell pseudo-bulk mixtures in linear scale]. a) pairwise Pearson correlation values between expression profiles for the different cell types, using a subset of the reference matrix containing only the markers used in the bulk deconvolution; b) pairwise Pearson correlation values between complete expression profiles for the different cell types; c) results using bulk deconvolution methods (nnls and CIBERSORT); d) results using single-cell deconvolution methods (MuSiC and DWLS). In c) and d), each grey column represents a specific cell type removed. Each data point conforming a boxplot represents a different scaling/normalization strategy used.

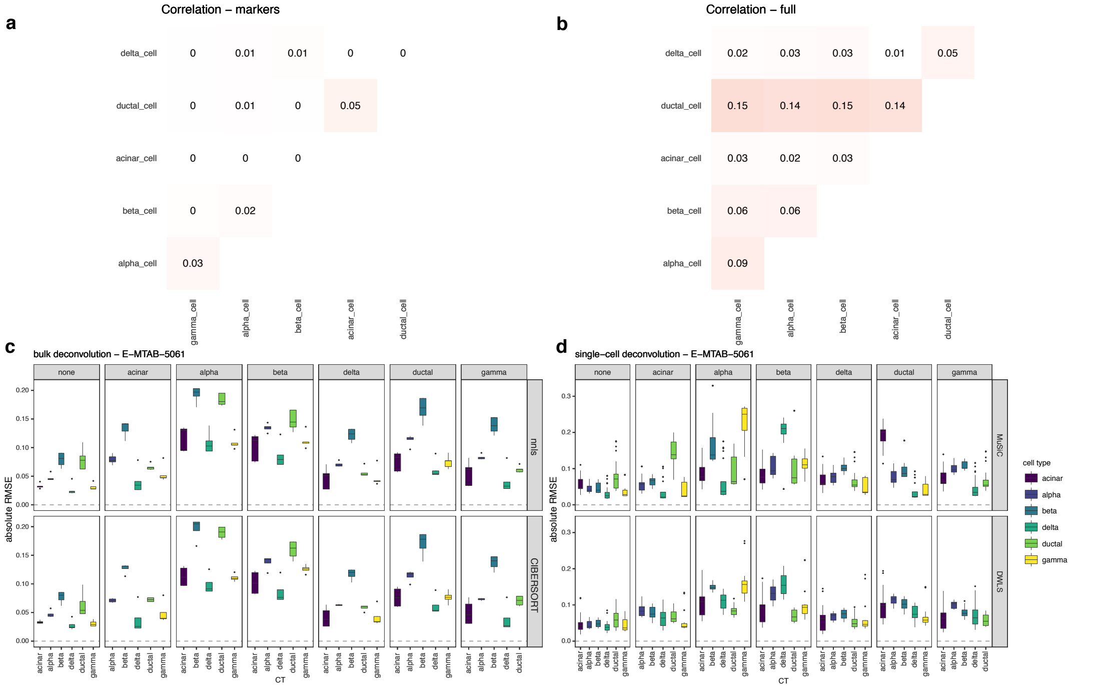


**Supplementary Figure 11** – Effect of cell type removal on the deconvolution results using the E-MTAB-5061 dataset [100-cell pseudo-bulk mixtures in linear scale]. a) pairwise Pearson correlation values between expression profiles for the different cell types, using a subset of the reference matrix containing only the markers used in the bulk deconvolution; b) pairwise Pearson correlation values between complete expression profiles for the different cell types; c) results using bulk deconvolution methods (nnls and CIBERSORT); d) results using single-cell deconvolution methods (MuSiC and DWLS). In c) and d), each grey column represents a specific cell type removed. Each data point conforming a boxplot represents a different scaling/normalization strategy used.

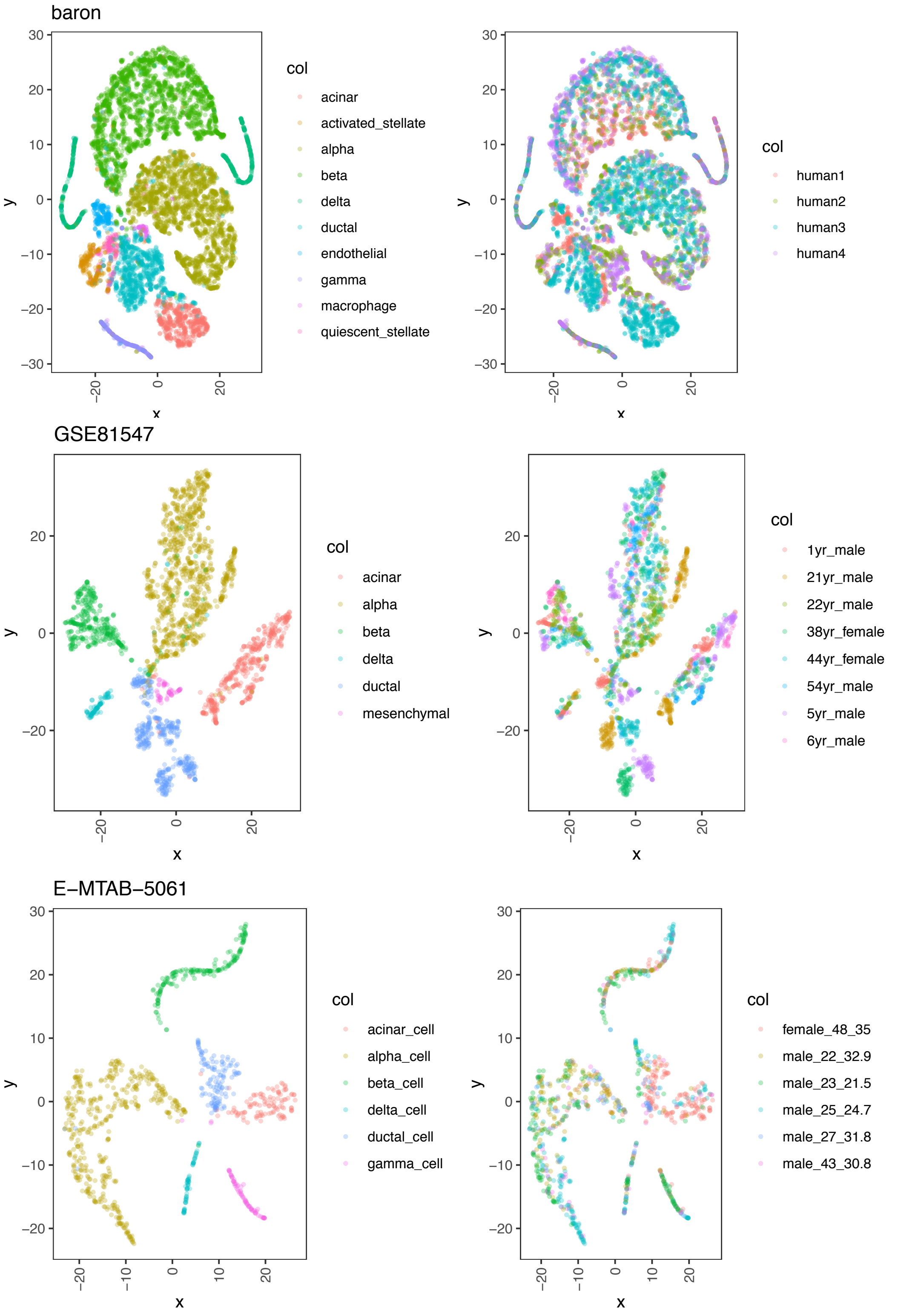


**Supplementary Figure 12** – Dimensionality reduction plots (tSNE) by cell type (left) and donor (right) across all datasets after quality control.
